# A novel fluorescent multi-domain protein construct reveals the individual steps of the unfoldase action of Hsp70

**DOI:** 10.1101/2022.02.17.480908

**Authors:** Satyam Tiwari, Bruno Fauvet, Salvatore Assenza, Paolo De los Rios, Pierre Goloubinoff

## Abstract

A detailed understanding of the mechanism by which Hsp70 chaperones protect cells against protein aggregation is hampered by the detailed characterization of the aggregates, which are typically heterogeneous. To tackle this problem, we designed here a reporter chaperone substrate, MLucV, composed of a stress-labile luciferase core, flanked by stress-resistant fluorescent mTFP and Venus domains, which upon denaturation formed a discrete stable population of small aggregates. Combining Förster Resonance Energy Transfer and enzymatic activity measurements provided unprecedent details on MLucV states, including native, aggregated, unfolded and chaperone-bound conformations. Using MLucV, we probed the various steps undertaken by bacterial Hsp70 to convert stable discrete aggregates into native proteins. The mechanism first involved an ATP-fuelled disaggregation and unfolding step of the stable pre-aggregated substrate, with a consequent stretching of MLucV beyond simply-unfolded conformations, followed, upon release, by native refolding. Furthermore, the ATP-fuelled unfolding action of Hsp70 on MLucV aggregates could accumulate native MLucV species under elevated denaturing temperatures, highly adverse to the native state. These results unambiguously excluded binding and preventing aggregation from the non-equilibirum mechanism by which Hsp70 converts stable aggregates into metastable native proteins.

## INTRODUCTION

Nascent polypeptides chains emerging from ribosomes can spontaneously acquire secondary and then near-native tertiary structures (Anfinsen 1973). Once released, newly translated proteins can either readily function, or translocate, and/or further assemble with other polypeptides to form functional oligomers (Wolynes, Onuchic et al. 1995, Dobson 2003). To carry out their unique biological function, native proteins need to remain relatively stable in the crowded environment of cells (Rivas and Minton 2016). Yet, exposure of native proteins to heat stress may affect their stability. Their partial unfolding may transiently expose hydrophobic residues to the aqueous phase, prompting them to seek more stable inactive misfolded and aggregated conformations (Ellis 2001). This is exemplified by the model reporter enzymes MDH (Goloubinoff, Sassi et al. 2018) and firefly luciferase (Sharma, De los Rios et al. 2010) that without chaperones and ATP become in a few minutes irreversibly inactivated at 37°C. The resulting aggregates come in various sizes and with several degrees of compactness, and when the heat stress is over, they do not revert to the soluble native state within a biologically-relevant time scale.

Misfolding and aggregation are intrinsic consequences of the physics of proteins and their cytotoxic effects is particularily affecting the neurons of complex metazoan (Hartl 2017). In cells, the Hsp70s, Hsp60s, Hsp90s and Hsp100s are distinct families of molecular chaperones that, assisted by a plethora of co-chaperones, use energy from ATP-hydrolysis to prevent and actively revert protein aggregation. Along the tree of life, this chaperone network has become progressively more complex concomitantly with an increase in the proteome complexity, in particular of the metazoans who became enriched with new misfolding- and aggregation-prone protein folds (Rebeaud, Mallik et al. 2021).

Strong evidence has accumulated that Hsp70s, Hsp100s and Hsp60s are acting as polypeptide unfolding enzymes that can specifically target stable misfolded and aggregated protein substrates, and use energy from ATP hydrolysis to convert them into unstable, unfolded monomeric conformations that, upon chaperone-release, may then spontaneously fold to the native state (Finka, Mattoo et al. 2016). Hsp70 functions in close collaboration with Hsp60s, Hsp90s and Hsp100s, but it can also act independently from them. As such, Hsp70 apparently serves as the central hub of the chaperone-based “protein repair” machineries in both prokaryotes and eukaryotes (Fauvet, Rebeaud et al. 2021, Rebeaud, Mallik et al. 2021).

The activity of Hsp70 strictly depends on a cohort of co-chaperones, called J-domain proteins (JDPs) for their common conserved J-domain that interacts with the ATP-bound conformation of Hsp70 (Kityk, Kopp et al. 2018). JDPs are in general multidomain proteins (Kampinga and Craig 2010) that, besides the J-domain, comprise other domains that can bind other macromolecules and macromolecular complexes, such as protein aggregates as in the case of DNAJAs and DNAJBs (Gillis, Schipper-Krom et al. 2013, Nillegoda, Stank et al. 2017), and also alternatively-folded protein oligomers, such as clathrin cages and the Heat Shock Factor 1 (Masser, Kang et al. 2019, Kmiecik, Le Breton et al. 2020). Other JDPs are associated to protein import pores across the ER and mitochondrial membranes (Frazier, Dudek et al. 2004, Jung and Kim 2021), thereby recruiting Hsp70 onto JDP-associated imported polypeptides. The targeted Hsp70 can in turn use energy from ATP hydrolysis to import proteins into organelles, and in general convert the JDP-associated macromolecular complexes into conformationally different ones, with different biological activities (Finka, Mattoo et al. 2016).

Upon the concurrent interaction of a J-domain and a bound polypeptide substrate with ATP-Hsp70, the ATPase cycle is accelerated up to a thousand-fold (Pierpaoli, Sandmeier et al. 1997, Han and Christen 2004), triggering Hsp70’s strong non-equilibrium binding to the misfolded substrate (ultra-affinity), that exploits the dramatic difference of the binding/unbinding rates between the ATP- and ADP-bound chaperone-substrate complexes (De Los Rios and Barducci 2014). While concomitantly inducing JDP-release, the ADP-Hsp70 induces substrate unfolding by clamping and entropic pulling (De Los Rios, Ben-Zvi et al. 2006, Hinault, Cuendet et al. 2010). Substrate release is then promoted by the Nucleotide Exchange Factors (NEF) (Brehmer, Gassler et al. 2004).

Crucial to a more precise elucidation of the mechanism of action of Hsp70 however, is the characterization of its aggregated protein substrates, a task that is complicated by the great heterogeneity of the non-native protein ensemble, comprising aggregates in many sizes, each with different degrees of compactness and solubility, as well as some misfolded monomers and transiently unfolded species (Diamant, Ben-Zvi et al. 2000).

To address this problem, we designed a model multidomain chaperone reporter protein: MLucV (Fig.S1). It is a monomeric, 120 kDa enzyme, composed of a heat-labile and urea-sensitive core made of a mutant active firefly luciferase (Sharma, De los Rios et al. 2010), flanked on either sides by urea- and thermo-resistant fluorescent domains. This allowed probing the various MLucV conformations by Förster Resonance Energy Transfer (FRET) and by luciferase enzymatic activity, both *in vitro* and in cells, finding that the various native, unfolded, chaperone-bound and aggregated states can be unambiguously distinguished. Furthermore, heat- and urea-pretreatments reproducibly formed small soluble MLucV aggregates, all within the narrow size range of ~12 polypeptides. The action of the Hsp70/JDP/NEF chaperone system (here bacterial DnaK/DnaJ/GrpE) could then be followed with unprecedented detail, revealing the various steps through which it disassembles stable protein aggregates, without necessitating assistance from the co-disaggregases ClpB. Hsp70 was further found to unfold and stretch the substrate beyond their urea-unfolded conformation. Moreover, the ATP-fuelled unfolding action of Hsp70 could actively disaggregate stable MLucV aggregates and accumulate native species against equilibrium, under stressful conditions that are highly adverse to the native state of the luciferase.

## RESULTS

### MLucV recapitulates the most important native and non-native states of proteins

The different MLucV states could be characterized by two independent and complementary means: FRET and enzymatic luciferase activity (see Figure S1 for the sequence of the construct). We first analyzed the FRET spectra of 400 nM native and MLucV in the presence of 4 M urea, as well as of an equimolar amount (400 nM each) of separated mTFP and Venus proteins, finding that they were clearly distinguishable from each other (Fig.1a). These spectra were converted into FRET efficiencies (see Methods), finding that the FRET efficiency of native, active MLucV (~0.43) was significantly higher than the baseline corresponding to the disconnected mTFP and Venus control (0.33) (Fig.1b; when making reference to relative FRET efficiencies, we normalized efficiencies so that 100% and 0% corresponded to the FRET of native MLucV and separated dyes respectively, as explained in the Methods section). This indicates that when connected by a compact native luciferase core, the two distal fluorophores are on average closer to each other than when they are unconnected. Denaturation of the luciferase core by increasing amounts of urea above 2.4 M completely abolished its enzymatic activity. Concomitantly, the FRET efficiency decreased down to ~0.35-0.36, a value that remained constant from 4-6 M and that was 73% lower than that of native MLucV, albeit significantly above the value of the unconnected mTFP and Venus control (Fig.1c). As a control, the fluorescence of individual Venus and mTFP did not change in the presence of up to 6 M urea (Fig.S2a,b), indicating that the FRET value of MLucV in the presence of 4 M urea was that of a species with a completely unfolded luciferase core, flanked by intact native fluorophores.

**Figure 1.**
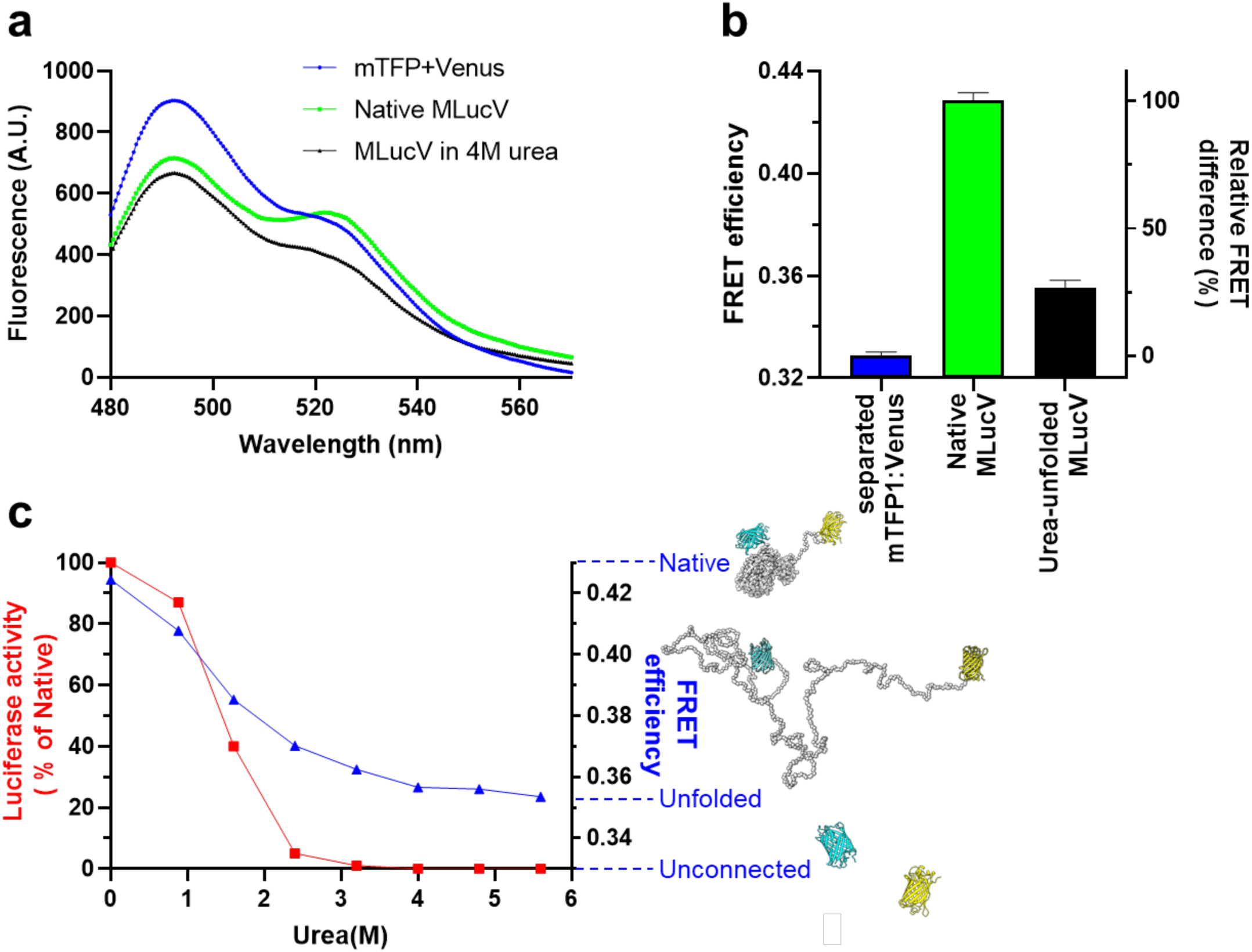
Biophysical characterization of native and non-native MLucV. **a**) FRET spectra of MLucV (0.4 *µ*M) under native conditions compared to 4M urea and to disconnected fluorophores (0.4 *µ*M each). **b**) Calculation of FRET efficiencies from the spectra in (a). The right y axis shows FRET efficiencies normalized to that of native MLucV (see Methods). **c**) MLucV Urea denaturation curves, monitored by FRET and luciferase activity. MLucV (0.4 μM) was incubated with different Urea concentrations (0, 0.5, 1, 1.5, 2, 3, 4, 6 M Urea) at 30°C for 10 min. Luciferase activity and FRET was measured in the presence of each urea concentration. The native and unfolded protein models have been obtained using Molecular Dynamics simulations (see Methods).

We next addressed the behavior of aggregated MLucV following two different denaturation protocols: urea pre-unfolding, followed by dilution, or plain heat denaturation. A short preincubation of 30 *µ*M MLucV in 4 M urea at 25°C, followed by a 75-fold dilution to 400 nM, readily produced luciferase species with a FRET spectrum that was distinct from those of native and urea-unfolded MLucV (Fig.2a compared to Fig.1a). This species was enzymatically inactive and had a FRET efficiency consistently higher by ~25% than that of the native species (Fig.2b compared with Fig.1b). This suggested that in the urea pre-unfolded species, mTFP and Venus were on average closer to each other than in the native MLucV monomers. Furthermore, no spontaneous refolding of urea pre-unfolded MLucV was observed, and its FRET remained stable for more than 90 minutes after initial denaturation and dilution (Fig.S3a,b). A similar result was obtained when 400 nM native MLucV was directly incubated for 23 minutes at 37.7°C: the corresponding fluorescence spectrum resembled that of urea-preunfolded & diluted MLucV species (Fig.2a), albeit with a slightly higher FRET efficiency (Fig.2b). Furthermore, over 98% of luciferase became inactive (Fig.2b) and remained so during two hours after the heat stress (Fig.S3b). In a control experiment, the intrinsic fluorescence of individual mTFP and Venus was found to be unaffected by a heat treatment of up to one hour at 39°C (Fig.S2c,d).

**Figure 2.**
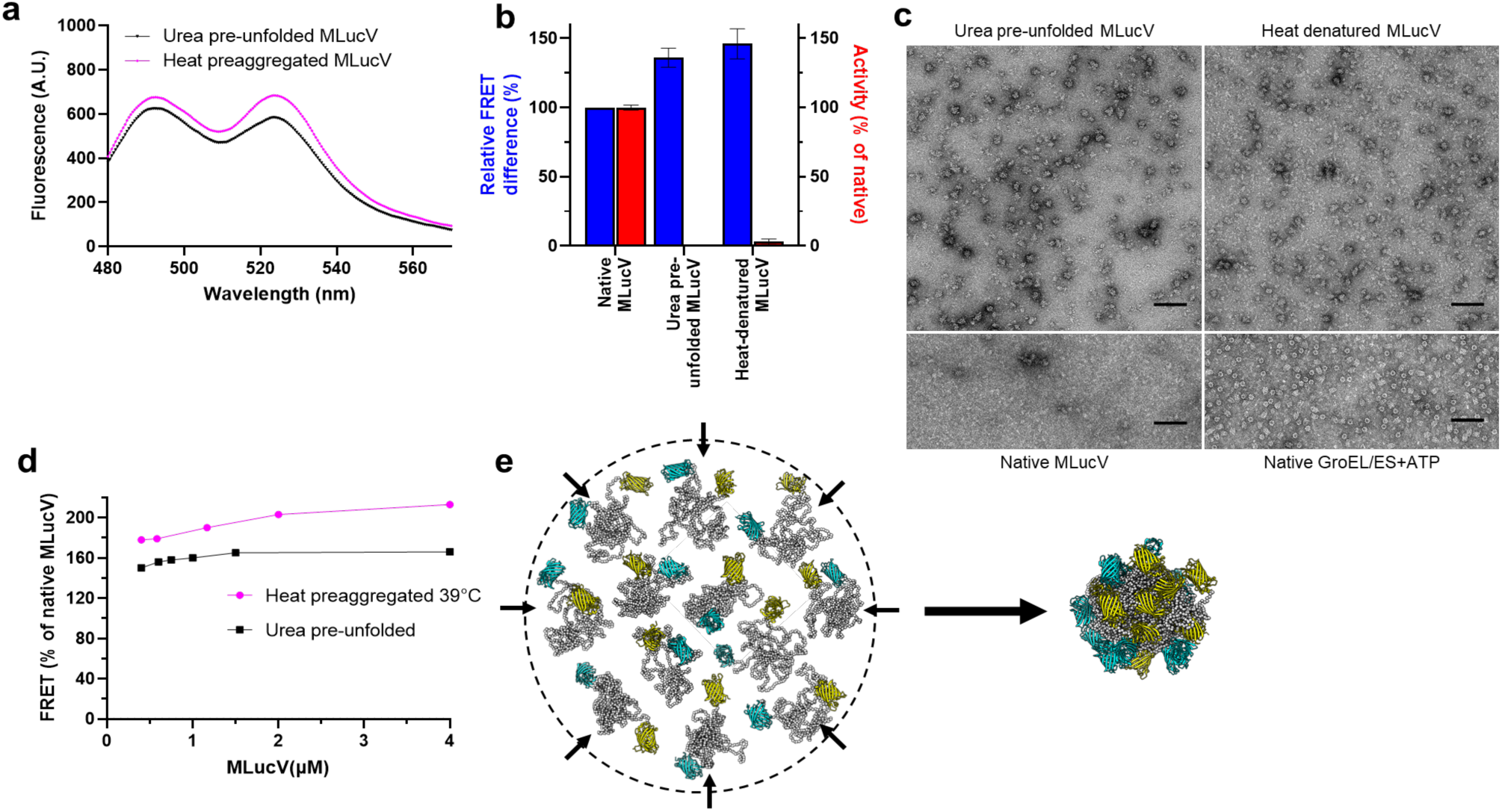
Biophysical characterization of aggregated MLucV. **a**) FRET spectra of 4 M urea pre-unfolded MLucV at 25°C and heat-preaggregated (23 min at 37.7°C) MLucV. **b**) FRET efficiencies derived from the spectra in (a) and luciferase activity in the corresponding conditions. The left y axis shows FRET efficiencies (blue bars) normalized to that of native MLucV, and the right y axis shows similarly normalized luciferase activity (red bars). **c**) The structure of urea pre-unfolded and heat-denatured MLucV analyzed by negative stain electron microscopy. Scale bar: 100 nm. Below: Native MLucV and native GroEL+GroES taken as controls. **d**) Relative FRET efficiency of MLucV complexes following 20 min preincubation at 39°C of MLucV, from 0.4 up to 4 μM, (magenta circles), or of 30 μM MLucV preunfolded at 25°C in 4 M urea, then diluted and incubated 20 min at 25°C (black squares). **e**) Molecular Dynamics simulation of the aggregation of 12 misfolded monomers: they were free to diffuse, rotate and rearrange in a confining potential (schematically represented by the circle), which became progressively more confining (arrows) until the 12 monomers arranged in a single aggregate. The confining potential was then turned off and the resulting aggregate allowed to briefly rearrange (see Methods).

Negative-stain electron microscopy showed that both urea pre-unfolded and heat-denatured MLucV species formed a population of discrete roughly spherical (although some were slightly elongated) particles with a diameter of about 22.3 ± 3.1 nm and 20.5 ± 3.3 nm, respectively (Fig.S4), which were distributed over a slightly wider range of sizes compared to the GroELS particles, used here as a control of complexes with a specific size (the observed average diameter of GroEL rings was 16.1 ± 1 nm). Particles obtained from urea pre-unfolding of MLucV were slightly larger, with a wider size distribution than the heat-denatured MLucV. Nevertheless, unprecedented for randomly generated protein aggregates, the size distribution of the MLucV aggregates remains very narrow relative to what usually occurs to aggregating proteins in general (Fig.2c and S4). Size Exclusion Chromatography – Right Angle Light Scattering (SEC-RALS) showed particles with an average mass of 1500 kDa, suggesting an average of 12 ± 2 misfolded MLucV subunits (Fig.S5a). In the case of urea pre-unfolded MLucV, the FRET efficiency of the aggregates as a function of protein concentration reached at 1500 nM a plateau of 166% of the native value, which was maintained when increasing the concentrations (by reducing the extent of dilution) up to 4000 nM of MLucV (Fig. 2d). Yet, here, 4000 nM MLucV formed aggregates similar in their FRET to those formed from a 1500 nM species. This was confirmed by SEC-RALS measurements at up to 2000 nM aggregating urea pre-unfolded MLucV, showing a similar size profile (Fig.S5b). These results are at variance with the general observation that increasing protein concentrations also increases the size and the compactness of protein aggregates (Meisl, Kirkegaard et al. 2016). Heat denaturation produced similar results, although the FRET efficiency values kept mildly increasing for all the tested concentrations of aggregating species (Fig.2d). Further experiments would be needed to characterize these oligomeric species and their dependence on the concentration of the misfolding species. Otherwise specified, throughout the remainder of this work, we used stable aggregates produced by pre-denaturing 30 micromolar MLucV for 5 minutes in 4 M urea, followed by a 75-fold dilution to a final concentration of 400 nM MLucV aggregated protomers.

Attesting for the solubility of these MLucV aggregates, the light-scattering signals for both urea- and heat-unfolded MLucV were very low and remained so over time, compared to an equimolar Luciferase control without flanking GFP domains (Fig.S6a,b). Moreover, following centrifugation, these inactive MLucV aggregated species remained soluble, at variance with aggregates of the Luciferase control (Fig.S6c).

We hypothesized that in a typical MLucV aggregate, the monomers should tend to arrange according to micellar principles, with their misfolded, insoluble luciferases sticking together in a core held together by hydrophobic interactions, and with the native, soluble hydrophilic, non-sticky mTFP donors and Venus acceptors mostly distributed at the surface. Their increased density on the particle’s surface would translate, as observed, in a higher FRET efficiency compared to that of the native monomers. Such a configuration for up to ~12 polypeptides would expectedly limit ensuing insertions of additional misfolded luciferases into the hydrophobic core, privileging the formation of more similarly-sized particles, over the further growth of the aggregates.

To address this idea, we performed simulations using a coarse-grained description of luciferase, with an inter-residue force-field tuned to describe non-native proteins, while the two soluble GFPs were maintained native by suitable structural restraints (see Methods). The simulation started with 12 MLucV misfolded monomers, namely polypeptides whose luciferase core was rather compact but not native. We then promoted their aggregation by introducing a confining spherical potential, whose radius shrinked in time. Yet, remarkably, no instructions were provided about the possible geometric arrangement of the luciferase and the GFPs in the final aggregate. Once the monomers were aggregated, the confining potential was switched off, and the system was let to further equilibrate. The simulation produced nearly spherical dodecameric particles, with most of the non-native luciferases packed inside and most of the GFPs exposed on the surface, positioned closer to each other than in the unfolded or the native monomeric states, thus supporting our model (Fig. 2e).

The limited size of these particles can be understood on simple geometric grounds: packing *n* MLucV monomers results in roughly spherical aggregates with a volume proportional to *n* and thus a radius proportional to *n*^1/3^. Correspondingly, the surface grows proportionally to *n*^2/3^, but must be covered by 2*n* GFPs. Thus, the number of GFPs (proportional to *n*) increases faster than the space they can occupy (proportional to *n*^2/3^), setting a limit to the number of monomers that can take part into a single aggregate particle. The precise value of this upper bound depends on several factors that are not fully characterized, such as the size and cohesive energy of the misfolded cores, and the fluctuation range of the soluble flanking domains.

The *in vitro* characterization of the different states of MLucV also allowed exploring its behavior *in vivo. E. coli* cells expressing low levels of MLucV at 30°C (to avoid the formation of inclusion bodies) displayed a fluorescence spectrum and a FRET efficiency for MLucV that was similar to those found *in vitro* for native MLucV (Fig.S7a and b, to be compared to Fig.1a and b). When the cells were incubated during 10 minutes at 39°C, the in-cell FRET efficiency increased and the luciferase activity from extracts correspondingly decreased (Fig.S7c), indicating that the luciferase lost its native-foldedness and the MLucV formed aggregates similar to those found *in vitro* (Fig.S7b, compared to Fig.2b) indicating that MLucV can be used as an *in vivo* reporter for the various stress-dependent states of a protein, in the presence of about 250 mg/ml of other proteins, including 42, 31, 22, 5 and 5 micromolar of GroEL, DnaK, Tig, ClpB and HtpG chaperones, respectively (Fauvet, Finka et al. 2021).

### Probing MLucV states during ATP-fuelled unfolding and disaggregation by bacterial Hsp70

The ATP-fuelled disaggregation mechanism of Hsp70 chaperones has been predominantly studied using ill-defined pre-aggregated model protein substrates, showing large and poorly reproducible distributions of oligomeric states, with variable degrees of compactness and solubility (Glover and Lindquist 1998, Goloubinoff, Mogk et al. 1999). Here, we could clearly and reproducibly identify the *native*, the *unfolded*, the *chaperone-stretched* and a uniquely *aggregated* states of MLucV, and thus we resolved, at least in part, the difficulties at investigating the ATP-fuelled action of the Hsp70 chaperone on otherwise very heterogeneous populations of misfolded substrates. We thus exploited the properties of MLucV to address the mechanism by which bacterial Hsp70 (DnaK), assisted by its DnaJ and GrpE cochaperones (the KJE system), can use the energy from ATP hydrolysis to convert stable, inactive, discrete, mostly dodecameric MLucV aggregates, into native, functional monomeric conformers. To this aim, MLucV (30 *µ*M) was first incubated in 4 M urea for 5 minutes (at 25°C), then diluted to 400 nM, without or with ATP, without or with KJE (4:1:2 *µ*M, respectively), and two hours later both FRET efficiency and luciferase activity were measured (Fig.3a). In the absence of KJE, pre-unfolded MLucV with ATP was enzymatically inactive and reached a FRET efficiency around 150%, compared to the native value (normalized to 100%), typical of the small aggregates described beforehand. The addition during urea-dilution of the KJE chaperone system, but without ATP, produced only a slightly lower FRET signal than that of MLucV aggregating alone, indicating an ineffective “holdase” activity of the chaperone. Conversely, in the presence of ATP, the KJE system promoted, as shown by FRET, a strong decompaction of the aggregates and the accumulation of native monomers, as assessed by enzymatic activity (Fig.3a).

**Figure 3:**
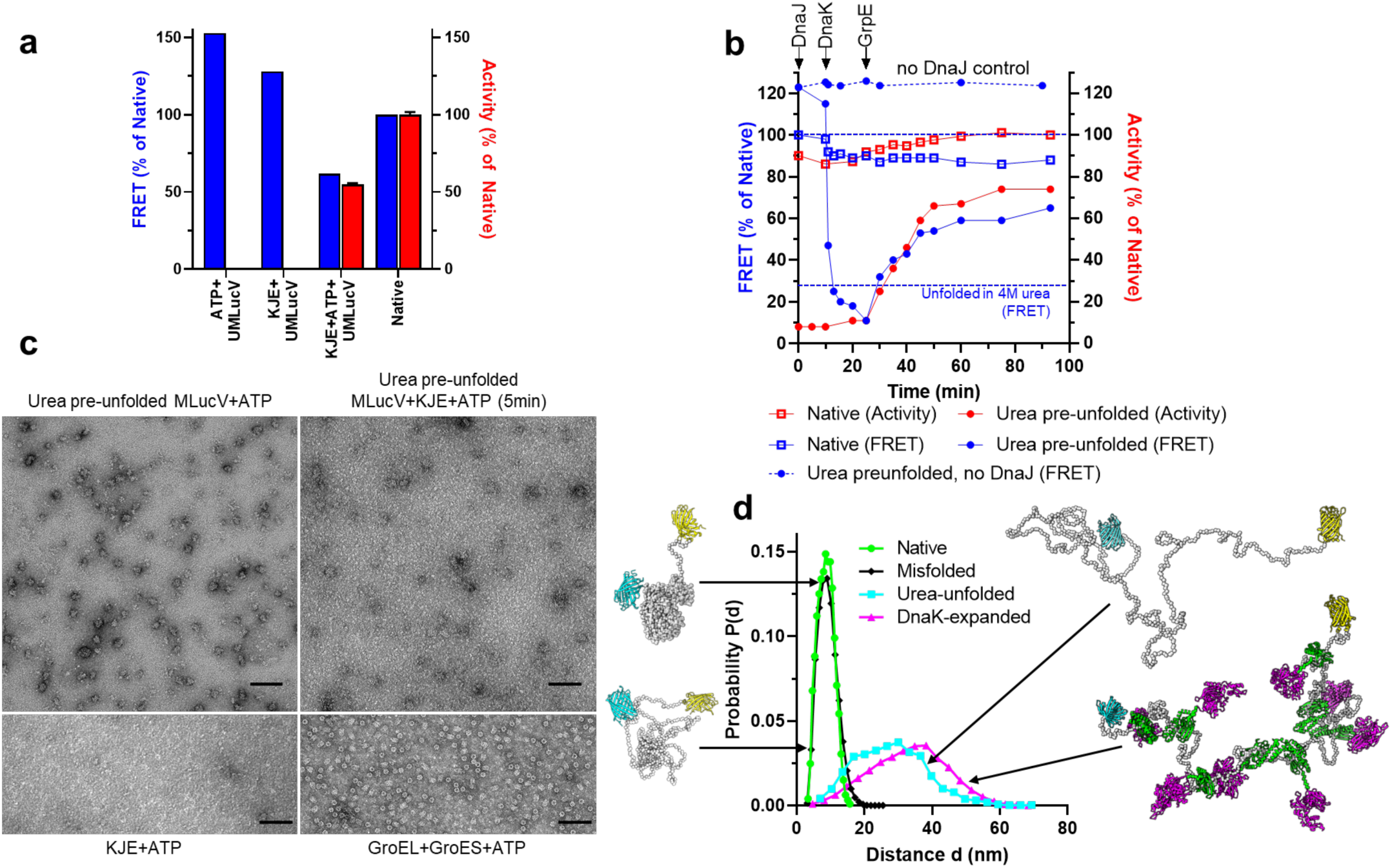
Resolving the individual steps of the KJE mechanism of action. **a**) 20 *µ*M MLucV pre-unfolded in 4 M urea at 25°C then diluted to 0.4 *µ*M in refolding buffer in the absence or presence of 4 mM ATP, in the absence or presence of KJE (4 *µ*M DnaK, 1 *µ*M DnaJ+2 *µ*M GrpE respectively). FRET and Luciferase activity was measured after 120 minutes at 25 °C. **b**) order-of-addition experiment showing DnaJ and DnaK-mediated unfolding of urea pre-unfolded MLucV in the presence of ATP, followed by GrpE-mediated release and refolding. FRET and luciferase activity were measured at 25°C; arrows indicate the time of addition of the KJE machinery components. Final concentrations of MLucV and chaperones were the same as in **a. c**) negative-staining electron micrographs of pre-unfolded MLucV (0.5 *µ*M) before (top left) and after (top right) a 5-minute incubation with the KJE+ATP chaperone system: DnaK (4 *µ*M), DnaJ (1 *µ*M) and GrpE (2 *µ*M) (top right panel). Bottom panels show control samples of KJE+ATP only (left) showing no large particles, and GroEL+GroES (2 *µ*M each, right). scale bar: 100nm. All samples were in the presence of 4 mM ATP. **d**) Probability distribution of the distance between the active centers of mTFP and Venus, collected from Molecular Dynamics simulations.

To address the effect of individual components in the reaction, we then applied an order-of-addition protocol. At the start of the assay, urea pre-unfolded inactive MLucV (0.4 μM protomers), in the presence of 4 mM ATP at 25°C, was first supplemented with DnaJ (1 μM) (Fig.3b). After ten minutes, only a minor ~5% decrease of the FRET efficiency was observed, without any increase of low basal native luciferase activity. This indicates that the expected binding of DnaJ on the surface of the aggregate had little to no effect on their compactness and on the aggregated luciferase core. In contrast, supplementing an eight fold excess of DnaK 10 minutes after DnaJ addition caused a dramatic and extensive decrease of the FRET efficiency, reaching a value lower than that attained by unfolded MLucV in the presence of 4 M urea. This very low FRET efficiency strongly suggests that most luciferase polypeptides, which were initially entangled within the core of the aggregated particles, have been stably bound by several DnaK molecules, up to eight on average, and must therefore have been solubilized, as evidenced by the observed greater distance between FRET pairs than in individual MLucV protomers. Indicating that most of the DnaK remained stably bound to the solubilized luciferase for at least 15 minutes and consistent with the extreme expansion of the luciferase core, no luciferase activity was recovered despite the presence of ATP and DnaJ. After 15 minutes, 2 μM of GrpE were added, which accelerated nucleotide exchange from ADP to ATP with the consequent release of the unfolded protein intermediate from DnaK. This led to a prompt recovery of native luciferase activity, up to ~70%, and a corresponding ~60% increase of the FRET efficiency towards the native values.

Remarkably, when this experiment was repeated without DnaJ (dotted line in Fig.3b), the addition of DnaK after 10 minutes did not change the high FRET efficiency of the MLucV aggregates and ultimately did not lead to any luciferase reactivation, despite the initial presence of ATP and the further later addition of GrpE. This clearly shows that although the aggregates are *bona fide* DnaK substrates, without DnaJ acting as the substrate-targeting co-chaperone, DnaK does not bind them.

The high specificity of the chaperone for substrate polypeptides that are misfolded and aggregated, and not for the native species, was further addressed by performing the same order of addition experiment with native MLucV (Fig.3b, open symbols). In the presence of ATP, addition of DnaJ to native MLucV caused no significant change in the FRET efficiency and in the apparently maximal luciferase activity (arbitrarily set in both cases to be 100%). The subsequent addition of DnaK after 10 minutes caused a sudden but minor ~9% drop of the FRET efficiency, whereas the luciferase activity remained high and essentially unchanged. The addition of GrpE after 25 minutes had no effect on the FRET efficiency, but led to the accumulation of ~ 10% additional active luciferase molecules. This suggests that the MLucV solution, which we initially considered to contain only native species, in fact likely contained up to 10% of inactive species with a higher FRET efficiency, suggesting they were aggregated. Nevertheless, this control experiment showed that even in the presence of ATP, neither DnaJ nor DnaK bound to the native MLucV conformers. The KJE system was able to specifically recognize the small minority of non-native MLucV polypeptides among a large excess of native conformers, attesting for the ability of this chaperone system to discriminate between misfolded conformations, acting as high-affinity substrates, and native conformations, acting as low-affinity products of the reaction. Noticeably, the FRET and activity values after 100 minutes did not reach 100%, (Fig.3a). Why is it so, whether by prolonged DnaK binding to MLucV, keeping it in a stretched conformation despite the presence of GrpE, possibly because of the increased ADP concentration, or just because of an excessively slow reactivation, which would lead to ~100% only on longer times, is left for future investigations.

The fast rate of chaperone-driven disaggregation and unfolding, as estimated from the FRET efficiency (~10% min^−1^), was confirmed by negative-stain electron microscopy. Within 5 minutes of KJE addition in the presence of ATP, about 55% of the particles disappeared, (from ~147 particles/*µ*m^2^ before KJE addition to ~67 particles/*µ*m^2^ after, Fig.3c). Moreover, the MLucV particles that were still present 5 minutes after KJE addition had distinctly fuzzier edges, suggesting that misfolded MLucV protomers at their surface were actively engaged in DnaK-mediated disaggregation and solubilization.

We then recapitulated the results obtained from the characterization of MLucV in its different states and from the unfolding action of DnaK using Molecular Dynamics simulations. By exploiting the same computational approach used above to investigate the organization of dodecamers (see Methods), we considered monomeric MLucV in four different conditions: i) native state; ii) unfolded (*i*.*e*. corresponding to the presence of a high concentration of Urea, see Methods); iii) misfolded state; iv) with eight ADP-DnaK molecules clamped onto the luciferase core. For each condition, the histogram of the distances between the mTFP and Venus active centers was collected (Fig.3d). The distance probability distribution of the native and misfolded monomers were very similar, suggesting that FRET alone could not tell the difference between the two, although the complementary measure of enzymatic activity could relieve this ambiguity. As we observed experimentally, the unfolding in 4 M Urea resulted in a much broader distance distribution, with an average distance that was larger than in the case of compact native and misfolded monomers, consequently leading to a much lower FRET efficiency, as experimentally observed. The simulated distance distribution was shifted towards even larger values for an MLucV monomer with 8 bound DnaK molecules, indicating that the binding of multiple chaperones could further stretch compact misfolded regions in a misfolded polypeptide, and consistent with the observation of a FRET efficiency even lower than that of urea-unfolded MLucV.

### Non-equilibrium activity of Hsp70

At 37.7°C, the native luciferase core of MLucV was highly unstable, losing its enzymatic activity at a rate of ~10% min^−1^ and reaching 83% inactivation in 30 minutes (Fig.4a). The subsequent addition of the full KJE chaperone system without ATP did not halt the subsequent denaturation of the luciferase. Yet, when alongside the added chaperones increasing concentrations of ATP were supplemented, native, active MLucV rapidly accumulated, at rate of 6% min^−1^, against its strong tendency to spontaneously denature at 37.7°C. The initial refolding rates were about the same for all the tested ATP concentrations, yet with 0.4 mM ATP, less than 50% of MLucV became transiently native and soon became inactive again, due to the consumption of the ATP. In contrast, with 6.4 mM ATP, a maximal non-equilibrium accumulation of the native population reached nearly 100% of the initial pre-denaturation level and it remained steadily high during more than one hour despite the strongly denaturing temperature. These results demonstrate that the KJE system can specifically target the low amounts of heat-misfolding species, as they keep forming at 37.7°C, while leaving untouched the conformers that are still native. At the same time, this also unambiguously demonstrates that the non-equilibrium stabilization of otherwise thermodynamically unstable native MLucV stringently depends on iterative, ATP-fueled chaperone unfolding cycles, which cease as soon as ATP is consumed.

**Figure 4:**
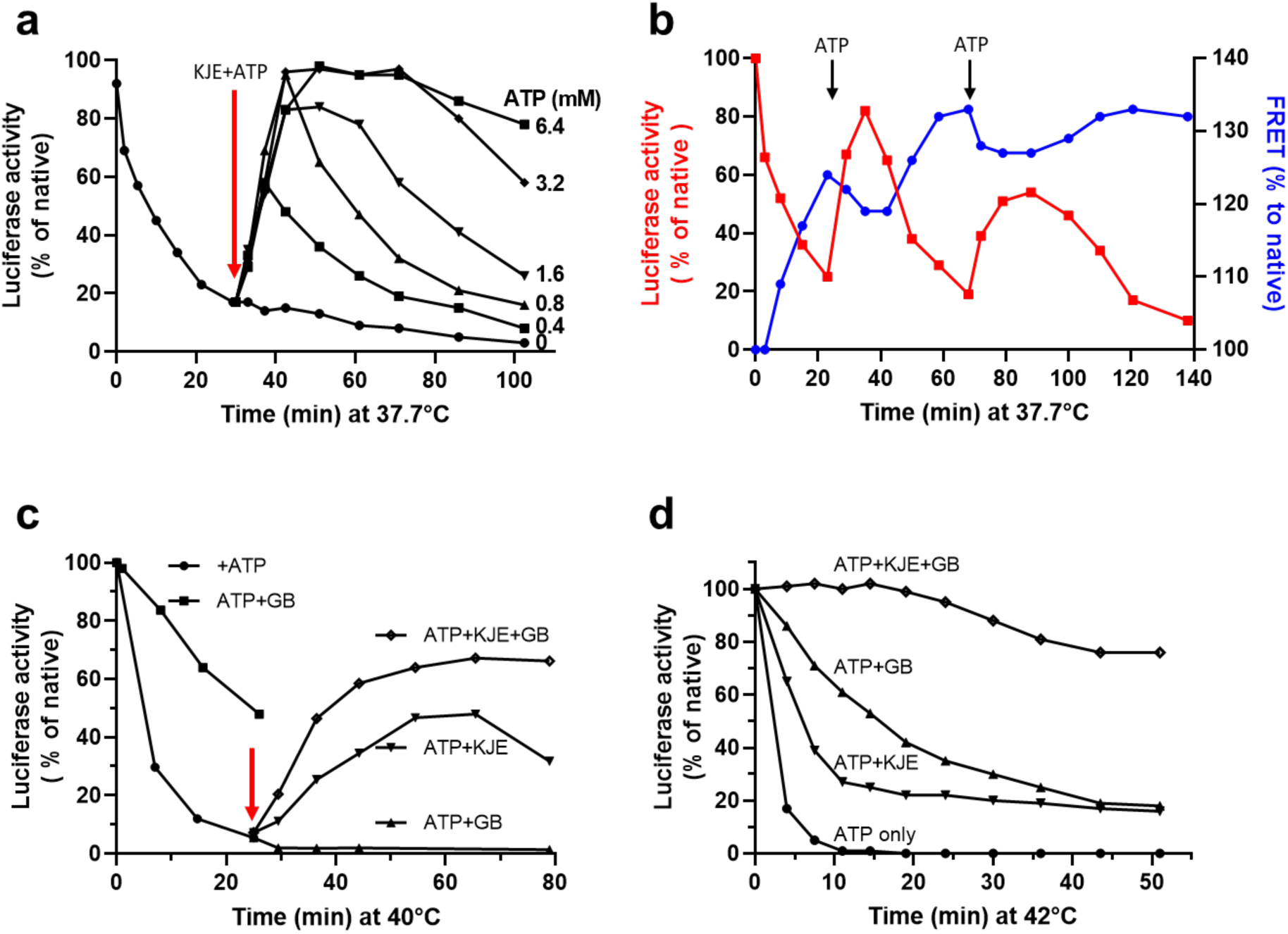
Non-equilibrium action of the KJE system. The KJE system can convert inactive species into native species under conditions that are unfavorable to the native state and this non-equilibrium process depends on ATP. **a**) 0.5 *µ*M native MLucV was incubated at 37.7°C in buffer+4 *µ*M Bovine Serum Albumin (BSA) for 30 min. At T=30 min KJE (4, 1, 2 *µ*M, respectively) and increasing amounts of ATP was added (red arrow) at the indicated concentrations and kept being incubated at 37.7°C. Luciferase activity is expressed in % of the maximal activity by native 0.5 *µ*M MLucV. **b**) 0.5 *µ*M native MLucV constantly incubated at 37.7°C, in the presence of 4 *µ*M BSA, 4 *µ*M DnaK, 1 *µ*M DnaJ, 2 *µ*M GrpE (KJE) and 2 *µ*M ClpB, first without ATP. At T=23 min, 0.8 mM ATP was added (first arrow), and again at T=68 min (second arrow). Luciferase activity and FRET efficiency were measured at the indicated time points and expressed as % of the initial luciferase activity and FRET efficiency at T=0. **c**) 0.4 *µ*M MLucV was incubated at 40°C in the presence of 5 mM ATP. At min 25 (red arrow), either 8% glycine betaine (GB), or KJE, or both were added. Luciferase activity was measured at the indicated time points. **d**) 0.5 *µ*M native MLucV was incubated at 42°C in the presence of 5 mM ATP or ATP+KJE or ATP+8% GB or both. Luciferase activity was measured at the indicated time points.

Working under heat-denaturing conditions further demonstrated that the chaperones cannot act by way of merely preventing the aggregation of their “client”, just by “holding” the unfolded or the already misfolded monomers. When native MLucV was first incubated at 37.7°C in the presence of the KJE system with the addition of the ClpB disaggregase (4μM DnaK, 1μM DnaJ, 2μM GrpE and 2μM ClpB), but without ATP, it lost its luciferase activity at an initial rate of ~10% min^−1^ (as in Fig.4a) and reached 25% activity in about 23 minutes (Fig.4b), as if chaperones were not present. Concomitantly, small aggregates were formed, as evidenced by the FRET signal that increased up to 125% of native MLucV, indicating that in the absence of ATP the protein-binding ability of a 2 fold excess of DnaJ, a 8 fold excess of DnaK and a 4 fold excess of ClpB was not effective at preventing MLucV aggregation. Addition, after 23 minutes, of a limiting amount of ATP (800 μM), caused a mild decrease of the FRET signal, indicating that disaggregation took place, despite the elevated temperature. Moreover, new native species were formed and rapidly accumulated during 12 minutes, reaching a maximal activity level of 82% of native MLucV, against their natural tendency to denature at 37.7°C. Yet, indicating that ATP was soon consumed, the luciferase activity dropped again and the FRET signal for aggregation concomitantly increased to a higher level than previously. This suggests that in the presence of ADP, none of the chaperones, although in 8 fold excess for DnaK alone, could passively “hold” the misfolded MLucV monomers and prevent their aggregation, contrary to the so-called “holdase” function generally ascribed to these chaperones (Winardhi, Tang et al. 2018, Hall 2020) Addition, after 68 minutes, of a second dose of 800 μM ATP further accumulated up to 46% native monomers in 32 minutes. Again, the eventual consumption of ATP led to a loss of the activity and to an increase in the FRET efficiency. Thus, without nucleotides or in the presence of ADP, none of the chaperones added to the solution were able to bind the misfolding species strongly enough to prevent their aggregation, whereas in the presence of ATP, the KJE chaperone could actively unfold non-native, misfolded species from the already formed aggregates and, upon release, allow them to transiently reach their native state. The prevention of aggregation activity was thus a mere by-product of the ATP-fuelled disaggregation, unfolding and refolding-upon-release activity of the KJE system, while an obligatory *holdase* step cannot explain the ATP-fuelled artificial reactivation of the substrate by the chaperones at high temperature.

The cytosol is a complex environment with high concentrations of various metabolites. We thus next addressed the ability of the metabolite glycine betaine (GB), which is known to accumulate in salt-stressed *E. coli* (Diamant, Rosenthal et al. 2003), to act alongside the KJE+ATP system at disassembling MLucV aggregates and recovering luciferase activity under a highly denaturing temperature (40°C). The presence at 40°C of 8% GB was found to slow down but, importantly not arrest, the denaturation of native MLucV, compared to the same treatment without GB, which instead caused the loss of 95% of the MLucV enzymatic activity after 25 minutes (Fig.4c). Addition of KJE+ATP at this time-point produced a slow accumulation up to ~ 50% of native MLucV, despite the ongoing heat-denaturing treatment at 40°C. Remarkably, addition of KJE+ATP together with GB lead to an even more rapid accumulation of up to 70% of native MLucV, indicating that under this heat-denaturing condition the partial thermo-protective action of GB on the native MLucV could also act in synergy with ATP-fuelled unfolding action of the chaperones. Adding KJE and GB to native MLucV at an even higher temperature, 42°C, was found to effectively maintain most of the luciferase activity, during 50 minutes of heat-shock, compared to the very rapid denaturation in their absence, and to the much milder effects of their individual presence (Fig.4d). Taken together, these results point to a complex interplay between chaperones and thermo-protective metabolites, modulating the efficacy of action of the KJE system in a non-trivial temperature-dependent way, likely by tuning the various rates of the folding/unfolding/misfolding/aggregation reaction to different degrees.

## DISCUSSION

The activity of proteins is often associated to their ability to switch between several distinct metastable native states (for example to exploit allosteric control for their function), and protein unfolding prior to degradation is a necessary step to recycle amino-acids for the synthesis of new proteins (Fauvet, Rebeaud et al. 2021). Evolution has thus often compromised between protein stability, necessary to withstand stresses, and conformational flexibility, necessary to carryout function. Therefore, the proteome of mesophilic and thermophilic organisms may still harbor a number of intrinsically thermolabile, or in general stress-sensitive proteins prone to lose their structure and activity and acquire instead more stable aggregated inactive conformations. ATPase chaperones represent an astute solution to this dilemma: through their non-equilibrium ATP-fuelled action they can increase the native population of metastable proteins without the need to increase the intrinsic thermodynamic stability of their native states, which would otherwise come at the cost of accrued rigidity. Thus, increasingly complex organisms along the tree of life, could have evolved an increasing number of very useful, albeit utterly chaperone-addicted, metastable proteins (Rebeaud et al., 2021). Nonetheless, the elucidation of the precise mechanism through which Hsp70 chaperones operate on non-native substrates has thus far been confused and complicated by the great heterogeneity of the non-native ensembles.

Here we designed a reporter protein, MLucV, that comprises a thermolabile luciferase core, and used it to probe the various states of a chaperone substrate while it is subject to ATP-driven protein disaggregation and unfolding by the bacterial Hsp70 system, under physiological and heat-stress conditions, both *in vitro* and *in vivo*. We showed that J-domain cochaperones target with exquisite specificity the Hsp70 molecules onto stress-misfolded and aggregated substrates, and not to native conformers. We showed that Hsp70s inject energy from ATP hydrolysis into the disassembly of stable aggregates and into the forceful unfolding of bound misfolded substrates, thus converting them into highly-stretched intermediates with a higher free energy, which upon release may spontaneously convert into more stable, low-affinity natively-refolded products.

Working under conditions where the native state of the MLucV luciferase core is intrinsically unstable (37.7°C), we further showed that in the absence of ATP, the mere “holdase” activity of the DnaK/DnaJ/GrpE system was ineffective at avoiding the formation of MLucV aggregates. Nonetheless, the binding of DnaJ to non-native polypeptides was necessary to recruit DnaK, which in its absence was unable to disassemble MLucV aggregates, even in the presence of ATP. Together, these observations suggest that the affinity of DnaJ for non-native monomers is high enough to target DnaK on them, but lower than the affinity that non-native monomers have for each other, thus ruling out DnaJ acting by way of its *holdase* function. Furthermore, no prevention of aggregation was observed by Hsp70 in the ADP-bound state, rebuking the usual view that ATP hydrolysis is needed to simply switch Hsp70 from the *low-affinity* ATP-bound state to the *high-affinity*, ADP-bound state. Instead, our results support the view of a truly non-equilibrium enhancement of affinity (De Los Rios and Barducci 2014) (*ultra-affinity*), exploiting the full ATPase cycle to increase the fraction of bound and consequently of the unfolded polypeptides intermediates, beyond what would be possible in either the ATP- or ADP-bound states. Last but not least, we also confirmed previous observations that the Hsp70 system is by itself a bona fide stand-alone disaggregation machinery (Diamant, Ben-Zvi et al. 2000). Our results stand in agreement with single-molecule FRET studies performed on small fluorophore-labeled luciferase at very low concentrations to avoid aggregation (Imamoglu, Balchin et al. 2020), while our usage of fluorescent proteins as FRET reporters also allows our reporter molecule to explore the action of chaperones on aggregates and to be used *in vivo* and in real time.

Reconstructing the early evolution of the chaperone network shows that following the two primordial chaperones present in the last common ancestor to all organisms (LUCA), namely an Hsp20 to prevent protein aggregations and ATP-fuelled Hsp60 to unfold small misfolded polypeptides, a Hsp70 system (DnaK, DnaJ and GrpE) disaggregase next appeared in the simplest terrestrial bacteria, and was soon to be acquired by primitive archea. Much later, only the more complex bacteria and the most complex archea evolved the Hsp70-dependent co-disaggregase ClpB, likely from a ClpCP-like protease, to render the basal stand-alone disaggregation by Hsp70 more efficient as observed in vitro (Diamant, Ben-Zvi et al. 2000). Only a billion year later, has Hsp110 emerged from Hsp70, to become a second Hsp70-dependent co-disaggregase in the cytosol and the ER of the first eukaryote, likely to increase the efficiency of disaggregation in organisms with more complex aggregation-prone proteomes (Rebeaud, Mallik et al. 2021).

## Materials and Methods

### MLucV (final reporter construct), mTFP1(FRET pair donor protein) and Venus (FRET pair acceptor protein) Plasmid construction

pTriEx-mTFP1(cp175)-Luciferase-Venus(cp173)-(called here MLucV) was created by using NEB Gibson assembly (NEBuilder HiFi DNA Assembly_E5520S) taking pTriEx mTFP1(cp175)-Barnase-Venus(cp173) as template plasmid (Fritz, Letzelter et al. 2013, Wood, Ormsby et al. 2018) in which Barnase was replaced by Luciferase (Sharma, De los Rios et al. 2010).

To construct a leaky expressing MLucV in E. coli,, MLucV was cloned into the low copy plasmid pSE380 (Genevaux et al., 2004). MLucV was PCR-amplified from the mTFP-ΔLuciferase-Venus-pTriEx4 plasmid as DNA template, using primers 5′-AGGAAACAGAATGGGCGGCCACCACCGC-3′ and 5′-CGCCAAAACATTAGATGTTGTGGCGGATCTTGAAGTTGG-3′. The pSE380 vector was PCR-amplified using primers 5’-CAACATCTAATGTTTTGGCGGATGAGAG-3’ and 5’-GGCCGCCCATTCTGTTTCCTGTGTGAAATTG-3’ and pSE380-MLucV was created using the NEB Gibson assembly kit (NEBuilder HiFi DNA Assembly_ E5520S). All constructs were confirmed by sanger sequencing.

GrpE and DnaJ were cloned into pET28-Smt3 expression vector. GrpE and DnaJ were first PCR amplified using E.coli genonic DNA then assembled by Gibson assembly (NEBuilder HiFi DNA Assembly_E5520S).

### Proteins

Recombinant production/purification: The MLucV construct was expressed in BL21-CodonPlus (DE3)-RIPL strain (Agilent) using ampicillin and chloramphenicol as the selection antibiotic. 20 mL of an overnight starter culture (grown in LB medium supplemented with 100 μg ml^−1^ ampicillin and 34 μg ml^−1^ chloramphenicol at 25 °C) was inoculated in 4000 mL of LB medium plus antibiotics and grown to an OD_600_ of 0.3 at 25 °C in a shaking incubator. The culture was cooled to 18°C (for 1 h, no shaking) and expression was induced with 0.2mM IPTG for overnight at 18 °C in a shaking incubator. Cells were harvested by centrifugation at 6000 r.p.m. for 12 min (pre-cooled rotor-JLA-9.1000 at 4°C, after that all steps at 4°C), and washed with chilled PBS. Cells were resuspended in 40 mL of lysis buffer (50mM Hepes-KOH pH 7.5, 300mM NaCl, 5mM EDTA, 5% glycerol, 5mM imidazole, 0.5 mM PMSF, 2 tablet of complete protease inhibitor cocktail (Roche) 2mM DTT, 40 mg total lysozyme powder(1mg/ml), and 200 μgDNase. Cells were disrupted by ultrasonication. The lysate was clarified by centrifugation at 15,000 r.p.m. for 30 min (JA 25.50 rotor, Beckman centrifuge) and applied directly onto a column of 2 ml pre-equilibrated Ni-NTA beads (cOmplete His-Tag Purification Resin from Merck). After several washes with high salt (500 mM NaCl), 20mM Imidazole and 5mM ATP, MLucV was eluted with 200mM imidazole in 25mM Hepes-KOH pH 7.5, 200 mM KCl, 10mM MgCl_2_, 2 mM DTT and 5% glycerol. Relatively pure fractions (purity of the fractions was assessed by 12% SDS-PAGE) were pooled, concentrated to ~5mg/ml (using 50kDa Amicon Ultra, Millipore), spin 10000 rpm 10 min, and loaded in superdex-200 increase gel filtration column (GE Healthcare), with final buffer (25 mM HEPES-KOH pH 7.5, 200 mM KCl, 10mM MgCl_2_, 2 mM DTT). Pure fractions were collected and stored in −80 °C.

DnaK of E.coli was expressed and purified as described previously (Feifel, Sandmeier et al. 1996). Purified DnaK was stored in 25 mM HEPES-KOH, 100 mM KCl, 10 mM MgCl2, pH 7.4, at −80 °C.

GrpE was expressed and purified from E. coli BL21 (DE3) cells harboring the pET28-Smt3-GrpE plasmid. Briefly, cells were grown in LB medium+kanamycin at 37 °C to OD_600_ ~0.4-0.5. Protein expression was induced by the addition of 0.5 mM IPTG for 3 hours. Cells were harvested, washed with chilled PBS and resuspended in buffer A (20 mM Tris-HCl pH 7.5, 200 mM NaCl, 5% glycerol, 2mM DTT, 20mM MgCl_2_) containing 5 mM imidazole, 1mg/ml Lysozyme, 1mM PMSF for 1 h. Cells were lysed by sonication. After high-speed centrifugation (16000 rpm, 30 min/4°C), the supernatant was loaded on to a gravity flow-based Ni-NTA metal affinity column (2 ml beads, cOmplete His-Tag Purification Resin from Merck), equilibrated and washed with 10 column volumes of buffer A containing 5 mM imidazole. After several washes with high salt, buffer A +200 mM NaCl, 20mM Imidazole and 5mM ATP, N-terminal His_10_-SUMO (small ubiquitin-related modifier) Smt3 tag was cleaved with Ulp1 protease (2mg/ml, 300 *µ*l, added to beads with buffer (20 mM Tris-HCl pH 7.5, 150mM KCl, 10mM MgCl2, 5% glycerol, 2mM DTT). Digestion of His_10_-Smt3 was performed on the Ni-NTA resin by, His_6_-Ulp1 protease. Because of dual His tags, His_6_-Ulp1 and His_10_-SUMO display a high affinity for Ni-NTA resin and remain bound to it during cleavage reaction. After overnight digestion at 4°C, the unbound fraction is collected (which contains only the native GrpE protein). GrpE was further purified by concentrating to ~3mg/ml and applying to a size-exclusion column (Superdex-200 increase, 10/30 GE heathcare) equilibrated in buffer A containing 5mM ATP. Pure fractions were pooled, concentrated by ultrafiltration using Amicon Ultra MWCO 10000 (Millipore), aliquoted and stored in −80 °C. DnaJ was purified in a similar way like GrpE.

All protein concentrations were determined spectrophotometrically at 562 nm using BCA Protein Assay Kit− Reducing Agent Compatible (cat no. 23250).

#### Control plasmids/proteins

Separated donor (mTFP1) and acceptor (Venus) fluorophores were prepared by PCR amplification from the pTriEx-MLucV plasmid using the Q5 Site-Directed Mutagenesis Kit (New England BioLabs). The proteins were expressed and purified in a similar way to MLucV. The plasmid pT7-lucC-His, carrying the luciferase gene of Photinus pyralis with the additional His6-coding sequence, was a gift from A.S. Spirin (Svetlov, Kolb et al. 2007). His-tagged Luciferase was purified as described previously (Sharma, De los Rios et al. 2010).

### FRET measurements and FRET efficiency calculation

All ensemble relative FRET efficiencies were calculated from maximum fluorescence emission intensities of donor (E_D_) and acceptor (E_A_) fluorophore by exiting donor only at 405 nm wavelength (Fritz, Letzelter et al. 2013, Wood, Ormsby et al. 2018). Fluorescence emission spectra analysis of MLucV reporter was performed on PerkinElmer LS55 fluorometer. Emission spectra were recorded from 480 to 580 nm wavelength with excitation slit 5 nm and emission slit 10nm. Average intensity values of spectral crosstalk was minimized by excitation donor at 405nm. Spectra were background-subtracted with spectra of buffer only or non-transformed cells in case of in-vivo measurements, samples acquired in the same conditions. The relative FRET efficiencies were calculated using the following equation:

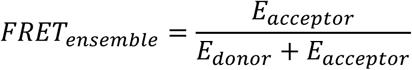

Normalized FRET efficiencies relative to that of native MLucV were calculated as follows:

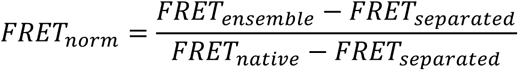

where FRET_ensemble_ is the measured ensemble FRET efficiency, FRET_separated_ is the calculated ensemble FRET measured in a solution of separated mTFP1 and Venus (0.33), and FRET_native_ is the measured ensemble FRET of native MLucV (0.43). Unless otherwise specified, all ensemble FRET measurements were performed at 400 nM of MLucV. Temperature was maintained at 25°C unless otherwise specified. All experiments were performed in LRB (20mM Hepes-KOH pH7.4,150mM KCl, 10mM MgCl_2_) refolding buffer containing 5 mM ATP, 2mM DTT, unless otherwise specified. 4*µ*M BSA was used in assays with chaperones to avoid MLucV species sticking to vessel, it has no effect on fate of the formed aggregates, nor have an effect on the activity of the chaperones. All experiments were repeated at least three times.

### FRET and activity analysis of live *E. coli* cells

10 mL of MLucV-expressing *E. coli* cells (strain W3110) were grown at 30°C in LB broth supplemented with antibiotics to OD_600_ ~ 0.4, 1mM IPTG was then added to induce the expression but only for 2 hours. Cells were pelleted and washed twice with PBS+100mM glucose pH 7.4 and resuspended in 10 mL of the same buffer. Equal number of cells in each sample was achieved by cell number normalization to OD_600_ = 1 by dilution. 100 μL of the cell suspension was added in cuvette with 100 μL buffer, mixing was performed by pipetting, and the fluorescence was recorded immediately at 25 °C or otherwise specified. To measure luciferase activity, same sample as for FRET, was sonicated 10 second with 2ug/ul lysozyme, immediately after lysis, 5*µ*l whole cells extract was taken to measure luciferase activity as published (Bischofberger, Han et al. 2003, Sharma, De los Rios et al. 2010) in a 96 well microtiter plate format. Four independent experiments were carried out with two technical replicates for each sample.

#### Luciferase activity assays

Luciferase activity was measured as described previously (Bischofberger, Han et al. 2003, Sharma, De los Rios et al. 2010). In the presence of oxygen, luciferase catalyzes the conversion of D-luciferin and ATP into oxyluciferin, CO_2_, AMP, PPi and hv. Generated photons were counted with a Victor Light 1420 Luminescence Counter from Perkin-Elmer in a 96 well microtiter plate format.

#### SEC-RALS analysis of Native MLucV, Urea pre-unfolded MLucV and Luciferase

Each sample (Native MLucV, Urea pre-unfolded MLucV or Luciferase, 1 *µ*M all samples) separately was loaded (300 *µ*l) in a superdex-200 increase column (GE Healthcare) attached to a Malvern Panalytical Omnisec resolve-reveal system. Using refractive index and right-angle light-scattering detectors, absolute molecular masses were calculated with OMNISEC-v11.10 software (Malvern Pananalytical) with the dn/dc value set to 0.185 mL/g, following calibration using the BSA monomer elution peak.

#### Light Scattering

To monitor aggregation propensity of urea denatured Luciferase and MLucV, 30 *µ*M Luciferase or MLucV was denatured with 4M urea at 25°C for 10 min, then diluted to a final concentration of 1 *µ*M in buffer A (50 mM Hepes-KOH pH7.5, 150 mM KCl, 10 mM MgCl_2_, 2 mM DTT), immediately aggregation was monitored by light scattering at 340 nm at 30°C for 30 min using Perkin Elmer Fluorescence Spectrophotometer. To monitor aggregation propensity of heat denatured Luciferase and MLucV, MLucV or Luciferase (0.5 *µ*M) was kept at 37.7°C in Buffer A having 4 *µ*M BSA and aggregation was monitored by light scattering at 340 nm for 30 min using Perkin Elmer Fluorescence Spectrophotometer.

#### Solubility test of urea pre-unfolded Luciferase and MLucV

30 *µ*M Luciferase or MLucV was denatured with 4M urea at 25°C for 10 min, then diluted to a final concentration of 1*µ*M in buffer (50 mM Hepes-KOH, pH 7.5 + 150 mM KCl + 10 mM MgCl_2_ + 2 mM DTT). Samples were incubated at 25°C for 30 min then soluble fraction was separated with high-speed centrifugation (14000 rpm, 10min). Equal volumes of Total and supernatant was loaded in 12% SDS-PAGE blue gel. For solubility of heat denatured MLucV, 0.5 *µ*M MLucV was incubated at 37.7°C for 30 min then soluble fraction was separated with high-speed centrifugation (14000 rpm, 10min). Equal volumes of Total and supernatant was loaded in 12% SDS-PAGE blue gel.

### Negative stain TEM Electron microscopy

Sample preparation for urea pre-unfolded MLucV species: 30 *µ*M MLucV was denatured with 4M urea at 25°C for 5 min, then diluted to a final concentration of 0.5 *µ*M in buffer (20 mM Hepes-KOH, pH 7.5 150 mM KCl +10 mM MgCl_2_ + 2 mM DTT + 5 mM ATP). Samples were incubated at 25°C for 30 min.

Sample preparation for heat denatured MLucV species-0.5 *µ*M MLucV was incubated at 37.7°C for 23 min in same buffer as for Urea preunfolded MLucV. Native control samples (MLucV and GroEL and GroES) were loaded directly on grid with same concentration as urea or heat denatured species.

For all above mentioned samples, 3 *µ*l sample (5 x diluted) was placed to freshly glow-discharged carbon EM grid (400 mesh copper, Electron Microscopy Sciences). After incubating for 2-minute, excess protein was removed by miliQ water wash, then grid was immediately placed on 100*µ*l droplet of 1% (w/v) uranyl acetate solution. After 1 min, excess uranyl acetate was removed from the grid by touching the edge with filter paper and grids were left to air-dry for 20 min. Images were acquired on a Philips CM100 Biotwin (80kV) transmission electron microscope. Micrographs were acquired at a magnification of 37,000X on an TVIPS F416 camera (4k x 4k). A minimum of 10 fields were screened per sample, to collect representative images, with 3 different repeats. The apparent particle sizes were quantified manually using the Fiji distribution of ImageJ, (Schindelin, Arganda-Carreras et al. 2012). At least 200 particles were measured per condition; because particles were elongated, two length measurements along perpendicular axes were taken, and the apparent size was determined as the average of the two. Normal distributions were then fitted to size histograms using MatLab R2019b.

### Molecular Dynamics simulations

We performed coarse-grained simulations following an approach already used previously (Assenza, Sassi et al. 2019). In short, all the amino acids were modeled as beads centered in the Cα atom. Native luciferase, mTFP and Venus were modelled as rigid bodies, and their native conformations obtained from the structures with pdb codes 1BA3 (Franks, Jenkins et al. 1998) 2HQK (Ai, Henderson et al. 2006) and 1MYW (Rekas, Alattia et al. 2002) respectively. In native conditions, linkers were modelled as disordered fragments by means of a model introduced previously (Smith, Ho et al. 2014) where we considered the case corresponding to the Monera hydrophobicity scale (Monera, Sereda et al. 1995). The extra amino acids at the C-terminus of the molecule used in our experiments were modelled as a disordered fragment. Specifically, the extra sequence was GGKSKLSYEQDGLHAGSPAALERAAA (see Fig.S1). Note that in the pdb entry 1BA3, the fragment GGKSKL is present in the fasta sequence, but absent in the pdb file; hence, it was considered here as part of the extra sequence. Under denaturing conditions, following the indications from experiments we assumed that mTFP and Venus kept their native structure, while luciferase was considered to be completely unfolded. In this case, the linkers and the unfolded luciferase were modelled by setting to zero the hydrophobicity of all the residues.

DnaK molecules in their ATP form were modelled by considering two independent rigid bodies for the Nucleotide-Binding Domain and the Substrate-Binding Domain (SBD), while the flexible linker was modelled as a disordered fragment. The rigid body of the SBD included also a bound heptapeptide. The reference structure was obtained by combining the pdbs 2KHO (Bertelsen, Chang et al. 2009) which provides the full structure of the ADP-bound DnaK, and 1DKX (Zhu, Zhao et al. 1996), which allows the modeling of the SBD bound to a heptapeptide. The combined structure was obtained by aligning the SBDs of both structures, as done in our previous work (Assenza, De Los Rios et al. 2015, Assenza, Sassi et al. 2019). The binding sites on luciferase were identified by means of a well-established algorithm (Rudiger, Germeroth et al. 1997), resulting in 13 non-overlapping binding sites.

Simulations were performed with LAMMPS (Thompson, Aktulga et al. 2022), with a Langevin thermostat set at temperature T=293 K and with damping parameter 16 ns^−1^. The timestep was set to 1 fs. In order to accelerate the convergence of simulations, each residue within a disordered fragment was assigned a mass equal to 1 Da, while residues belonging to rigid bodies were given a mass equal to 0.01 Da. Simulations were run for 2×10^7^ timesteps. In order to ensure full equilibration, the first 10^6^ steps were discarded from the analysis. Five independent simulations were performed for each case.

## Acknowledgements

This work was supported by grant 31003A_175453 from the Swiss National Fund. We thank Tatiana Mbefo Kamdem for technical assistance with protein purification.

## Author contributions

P.G. and P.D.L.R. supervised the project, S.T., P.D.L.R, and P.G. designed the experiments, S.T. generated reagents (plasmids and proteins), S.T. and P.G. performed the experiments, S.A. performed molecular dynamics simulations, S.T., P.G., S.A. and B.F. analyzed the data and created the figures. S.T., S.A., B.F, P.G., and P.D.L.R. wrote the manuscript. All authors read, edited and approved the manuscript.

## References

Ai, H. W., J. N. Henderson, S. J. Remington and R. E. Campbell (2006). “Directed evolution of a monomeric, bright and photostable version of Clavularia cyan fluorescent protein: structural characterization and applications in fluorescence imaging.” Biochem J 400(3): 531–540.

Anfinsen, C. B. (1973). “Principles that govern the folding of protein chains.” Science 181(4096): 223–230.

Assenza, S., P. De Los Rios and A. Barducci (2015). “Quantifying the role of chaperones in protein translocation by computational modeling.” Front Mol Biosci 2: 8.

Assenza, S., A. S. Sassi, R. Kellner, B. Schuler, P. De Los Rios and A. Barducci (2019). “Efficient conversion of chemical energy into mechanical work by Hsp70 chaperones.” Elife 8.

Bertelsen, E. B., L. Chang, J. E. Gestwicki and E. R. Zuiderweg (2009). “Solution conformation of wild-type E. coli Hsp70 (DnaK) chaperone complexed with ADP and substrate.” Proc Natl Acad Sci U S A 106(21): 8471–8476.

Bischofberger, P., W. Han, B. Feifel, H. J. Schonfeld and P. Christen (2003). “D-Peptides as inhibitors of the DnaK/DnaJ/GrpE chaperone system.” J Biol Chem 278(21): 19044–19047.

Brehmer, D., C. Gassler, W. Rist, M. P. Mayer and B. Bukau (2004). “Influence of GrpE on DnaK-substrate interactions.” J Biol Chem 279(27): 27957–27964.

De Los Rios, P. and A. Barducci (2014). “Hsp70 chaperones are non-equilibrium machines that achieve ultra-affinity by energy consumption.” Elife 3: e02218.

De Los Rios, P., A. Ben-Zvi, O. Slutsky, A. Azem and P. Goloubinoff (2006). “Hsp70 chaperones accelerate protein translocation and the unfolding of stable protein aggregates by entropic pulling.” Proc Natl Acad Sci U S A 103(16): 6166–6171.

Diamant, S., A. P. Ben-Zvi, B. Bukau and P. Goloubinoff (2000). “Size-dependent disaggregation of stable protein aggregates by the DnaK chaperone machinery.” J Biol Chem 275(28): 21107–21113.

Diamant, S., D. Rosenthal, A. Azem, N. Eliahu, A. P. Ben-Zvi and P. Goloubinoff (2003). “Dicarboxylic amino acids and glycine-betaine regulate chaperone-mediated protein-disaggregation under stress.” Mol Microbiol 49(2): 401–410.

Dobson, C. M. (2003). “Protein folding and misfolding.” Nature 426(6968): 884–890.

Ellis, R. J. (2001). “Macromolecular crowding: an important but neglected aspect of the intracellular environment.” Curr Opin Struct Biol 11(1): 114–119.

Fauvet, B., A. Finka, M. P. Castanie-Cornet, A. M. Cirinesi, P. Genevaux, M. Quadroni and P. Goloubinoff (2021). “Bacterial Hsp90 Facilitates the Degradation of Aggregation-Prone Hsp70-Hsp40 Substrates.” Front Mol Biosci 8: 653073.

Fauvet, B., M. E. Rebeaud, S. Tiwari, P. De Los Rios and P. Goloubinoff (2021). “Repair or Degrade: the Thermodynamic Dilemma of Cellular Protein Quality-Control.” Front Mol Biosci 8: 768888.

Feifel, B., E. Sandmeier, H. J. Schonfeld and P. Christen (1996). “Potassium ions and the molecular-chaperone activity of DnaK.” Eur J Biochem 237(1): 318–321.

Finka, A., R. U. Mattoo and P. Goloubinoff (2016). “Experimental Milestones in the Discovery of Molecular Chaperones as Polypeptide Unfolding Enzymes.” Annu Rev Biochem 85: 715–742.

Franks, N. P., A. Jenkins, E. Conti, W. R. Lieb and P. Brick (1998). “Structural basis for the inhibition of firefly luciferase by a general anesthetic.” Biophys J 75(5): 2205–2211.

Frazier, A. E., J. Dudek, B. Guiard, W. Voos, Y. Li, M. Lind, C. Meisinger, A. Geissler, A. Sickmann, H. E. Meyer, V. Bilanchone, M. G. Cumsky, K. N. Truscott, N. Pfanner and P. Rehling (2004). “Pam16 has an essential role in the mitochondrial protein import motor.” Nat Struct Mol Biol 11(3): 226–233.

Fritz, R. D., M. Letzelter, A. Reimann, K. Martin, L. Fusco, L. Ritsma, B. Ponsioen, E. Fluri, S. Schulte-Merker, J. van Rheenen and O. Pertz (2013). “A versatile toolkit to produce sensitive FRET biosensors to visualize signaling in time and space.” Sci Signal 6(285): rs12.

Gillis, J., S. Schipper-Krom, K. Juenemann, A. Gruber, S. Coolen, R. van den Nieuwendijk, H. van Veen, H. Overkleeft, J. Goedhart, H. H. Kampinga and E. A. Reits (2013). “The DNAJB6 and DNAJB8 protein chaperones prevent intracellular aggregation of polyglutamine peptides.” J Biol Chem 288(24): 17225–17237.

Glover, J. R. and S. Lindquist (1998). “Hsp104, Hsp70, and Hsp40: a novel chaperone system that rescues previously aggregated proteins.” Cell 94(1): 73–82.

Goloubinoff, P., A. Mogk, A. P. Zvi, T. Tomoyasu and B. Bukau (1999). “Sequential mechanism of solubilization and refolding of stable protein aggregates by a bichaperone network.” Proc Natl Acad Sci U S A 96(24): 13732–13737.

Goloubinoff, P., A. S. Sassi, B. Fauvet, A. Barducci and P. De Los Rios (2018). “Chaperones convert the energy from ATP into the nonequilibrium stabilization of native proteins.” Nat Chem Biol 14(4): 388–395.

Hall, D. (2020). “On the nature of the optimal form of the holdase-type chaperone stress response.” FEBS Lett 594(1): 43–66.

Han, W. and P. Christen (2004). “cis-Effect of DnaJ on DnaK in ternary complexes with chimeric DnaK/DnaJ-binding peptides.” FEBS Lett 563(1-3): 146–150.

Hartl, F. U. (2017). “Protein Misfolding Diseases.” Annu Rev Biochem 86: 21–26.

Hinault, M. P., A. F. Cuendet, R. U. Mattoo, M. Mensi, G. Dietler, H. A. Lashuel and P. Goloubinoff (2010). “Stable alpha-synuclein oligomers strongly inhibit chaperone activity of the Hsp70 system by weak interactions with J-domain co-chaperones.” J Biol Chem 285(49): 38173–38182.

Imamoglu, R., D. Balchin, M. Hayer-Hartl and F. U. Hartl (2020). “Bacterial Hsp70 resolves misfolded states and accelerates productive folding of a multi-domain protein.” Nat Commun 11(1): 365.

Jung, S. J. and H. Kim (2021). “Emerging View on the Molecular Functions of Sec62 and Sec63 in Protein Translocation.” Int J Mol Sci 22(23).

Kampinga, H. H. and E. A. Craig (2010). “The HSP70 chaperone machinery: J proteins as drivers of functional specificity.” Nat Rev Mol Cell Biol 11(8): 579–592.

Kityk, R., J. Kopp and M. P. Mayer (2018). “Molecular Mechanism of J-Domain-Triggered ATP Hydrolysis by Hsp70 Chaperones.” Mol Cell 69(2): 227–237 e224.

Kmiecik, S. W., L. Le Breton and M. P. Mayer (2020). “Feedback regulation of heat shock factor 1 (Hsf1) activity by Hsp70-mediated trimer unzipping and dissociation from DNA.” EMBO J 39(14): e104096.

Masser, A. E., W. Kang, J. Roy, J. Mohanakrishnan Kaimal, J. Quintana-Cordero, M. R. Friedlander and C. Andreasson (2019). “Cytoplasmic protein misfolding titrates Hsp70 to activate nuclear Hsf1.” Elife 8.

Meisl, G., J. B. Kirkegaard, P. Arosio, T. C. Michaels, M. Vendruscolo, C. M. Dobson, S. Linse and T. P. Knowles (2016). “Molecular mechanisms of protein aggregation from global fitting of kinetic models.” Nat Protoc 11(2): 252–272.

Monera, O. D., T. J. Sereda, N. E. Zhou, C. M. Kay and R. S. Hodges (1995). “Relationship of sidechain hydrophobicity and alpha-helical propensity on the stability of the single-stranded amphipathic alpha-helix.” J Pept Sci 1(5): 319–329.

Nillegoda, N. B., A. Stank, D. Malinverni, N. Alberts, A. Szlachcic, A. Barducci, P. De Los Rios, R. C. Wade and B. Bukau (2017). “Evolution of an intricate J-protein network driving protein disaggregation in eukaryotes.” Elife 6.

Pierpaoli, E. V., E. Sandmeier, A. Baici, H. J. Schonfeld, S. Gisler and P. Christen (1997). “The power stroke of the DnaK/DnaJ/GrpE molecular chaperone system.” J Mol Biol 269(5): 757–768.

Rebeaud, M. E., S. Mallik, P. Goloubinoff and D. S. Tawfik (2021). “On the evolution of chaperones and cochaperones and the expansion of proteomes across the Tree of Life.” Proc Natl Acad Sci U S A 118(21).

Rekas, A., J. R. Alattia, T. Nagai, A. Miyawaki and M. Ikura (2002). “Crystal structure of venus, a yellow fluorescent protein with improved maturation and reduced environmental sensitivity.” J Biol Chem 277(52): 50573–50578.

Rivas, G. and A. P. Minton (2016). “Macromolecular Crowding In Vitro, In Vivo, and In Between.” Trends Biochem Sci 41(11): 970–981.

Rudiger, S., L. Germeroth, J. Schneider-Mergener and B. Bukau (1997). “Substrate specificity of the DnaK chaperone determined by screening cellulose-bound peptide libraries.” EMBO J 16(7): 1501–1507.

Schindelin, J., I. Arganda-Carreras, E. Frise, V. Kaynig, M. Longair, T. Pietzsch, S. Preibisch, C. Rueden, S. Saalfeld, B. Schmid, J. Y. Tinevez, D. J. White, V. Hartenstein, K. Eliceiri, P. Tomancak and A. Cardona (2012). “Fiji: an open-source platform for biological-image analysis.” Nat Methods 9(7): 676–682.

Sharma, S. K., P. De los Rios, P. Christen, A. Lustig and P. Goloubinoff (2010). “The kinetic parameters and energy cost of the Hsp70 chaperone as a polypeptide unfoldase.” Nat Chem Biol 6(12): 914–920.

Smith, W. W., P. Y. Ho and C. S. O’Hern (2014). “Calibrated Langevin-dynamics simulations of intrinsically disordered proteins.” Phys Rev E Stat Nonlin Soft Matter Phys 90(4): 042709.

Svetlov, M. S., V. A. Kolb and A. S. Spirin (2007). “[Folding of the firefly luciferase polypeptide chain with immobilized C-terminus].” Mol Biol (Mosk) 41(1): 96–102.

Thompson, A. P., H. M. Aktulga, R. Berger, D. S. Bolintineanu, W. M. Brown, P. S. Crozier, P. J. in ‘t Veld, A. Kohlmeyer, S. G. Moore, T. D. Nguyen, R. Shan, M. J. Stevens, J. Tranchida, C. Trott and S. J. Plimpton (2022). “LAMMPS - a flexible simulation tool for particle-based materials modeling at the atomic, meso, and continuum scales.” Computer Physics Communications 271: 108171.

Winardhi, R. S., Q. Tang, H. You, M. Sheetz and J. Yan (2018). “The holdase function of <em>Escherichia coli</em> Hsp70 (DnaK) chaperone.” bioRxiv: 05854.

Wolynes, P. G., J. N. Onuchic and D. Thirumalai (1995). “Navigating the folding routes.” Science 267(5204): 1619–1620.

Wood, R. J., A. R. Ormsby, M. Radwan, D. Cox, A. Sharma, T. Vopel, S. Ebbinghaus, M. Oliveberg, G. E. Reid, A. Dickson and D. M. Hatters (2018). “A biosensor-based framework to measure latent proteostasis capacity.” Nat Commun 9(1): 287.

Zhu, X., X. Zhao, W. F. Burkholder, A. Gragerov, C. M. Ogata, M. E. Gottesman and W. A. Hendrickson (1996). “Structural analysis of substrate binding by the molecular chaperone DnaK.” Science 272(5268): 1606–1614.

