## Supplementary Figures for "A novel fluorescent multi-domain protein construct reveals the individual steps of the unfoldase action of Hsp70"

**a**

MAHHHHHHGSGEQKLISEEDLGSGSGSGGHHRVDFKTIYRAKKAVKLPDYHFVDHRIEILN
HDKDYNKVTVYESAVARNSTDGMDELYKGASGGMVSKGEETTMGVIKPDMKIKLKMEGNVN
GHAFVIEGEGEGKPYDGTNTINLEVKEGAPLPFSYDILTTAFAYGNRAFTKYPDDIPNYFK
QSFPEGYSWERTMTFEDKGIVKVKSDISMEEDSFIYEIHLKGENFPPNGPVMQKKTTGWDA
STERMYVRDGVLKGDVKHKLLLEGSGMEDAKNIKKGPAPFYPLEDGTAGEQLHKAMKRYAL
VPGTIAFTDAHIEVNITYAEYFEMSVRLAEAMKRYGLNTNHRIVVCSENSLQFFMPVLGAL
FIGVAVAPANDIYNERELLNSMNISQPTVVFVSKKGLQKILNVQKKLPIIQKIIIMDSKTD
YQGFQSMYTFVTSHLPPGFNEYDFVPESFDRDKTIALIMNSSGSTGLPKGVALPHRTACVR
FSHARDPIFGNQIIPDTAILSVVPFHHGFGMFTTLGYLICGFRVVLMYRFEEELFLRSLQD
YKIQSALLVPTLFSFFAKSTLIDKYDLSNLHEIASGGAPLSKEVGEAVAKRFHLPGIRQGY
GLTETTSAILITPEGDDKPGAVGKVVPFFEAKVVDLDTGKTLGVNQRGELCVRGPMIMSGY
VNNPEATNALIDKDGWLHSGDIAYWDEDEHFFIVDRLKSLIKYKGYQVAPAELESILLQHP
NIFDAGVAGLPDDDAGELPAAVVVLEHGKTMTEKEIVDYVASQVTTAKKLRGGVVFVDEVP
KGLTGKLDARKIREILIKAKKGGKSKLSYEQDGLHAGSPAALERAAAMDGGVQLADHYQQN
TPIGDGPVLLPDNHYLSYQSALSKDPNEKRDHMVLLEFVTAAGITLGMDELYKGGSGGMVS
KGEELFTGVVPILVELDGDVNGHKFSVSGEGEGDATYGKLTLKLICTTGKLPVPWPTLVTT
LGYGLMCFARYPDHMKQHDFFKSAMPEGYVQERTIFFKDDGNYKTRAEVKFEGDTLVNRIE
LKGIDFKEDGNILGHKLEYNYNSHNVYITADKQKNGIKANFKIRHNIEGTDILQKKLEELE
LDE

**Figure S1**: Amino acid sequence of MLucV (mTFP1-Luciferase-Venus). Cyan: the donor- mTFP, Yellow: the acceptor- Venus, magenta: Luciferase

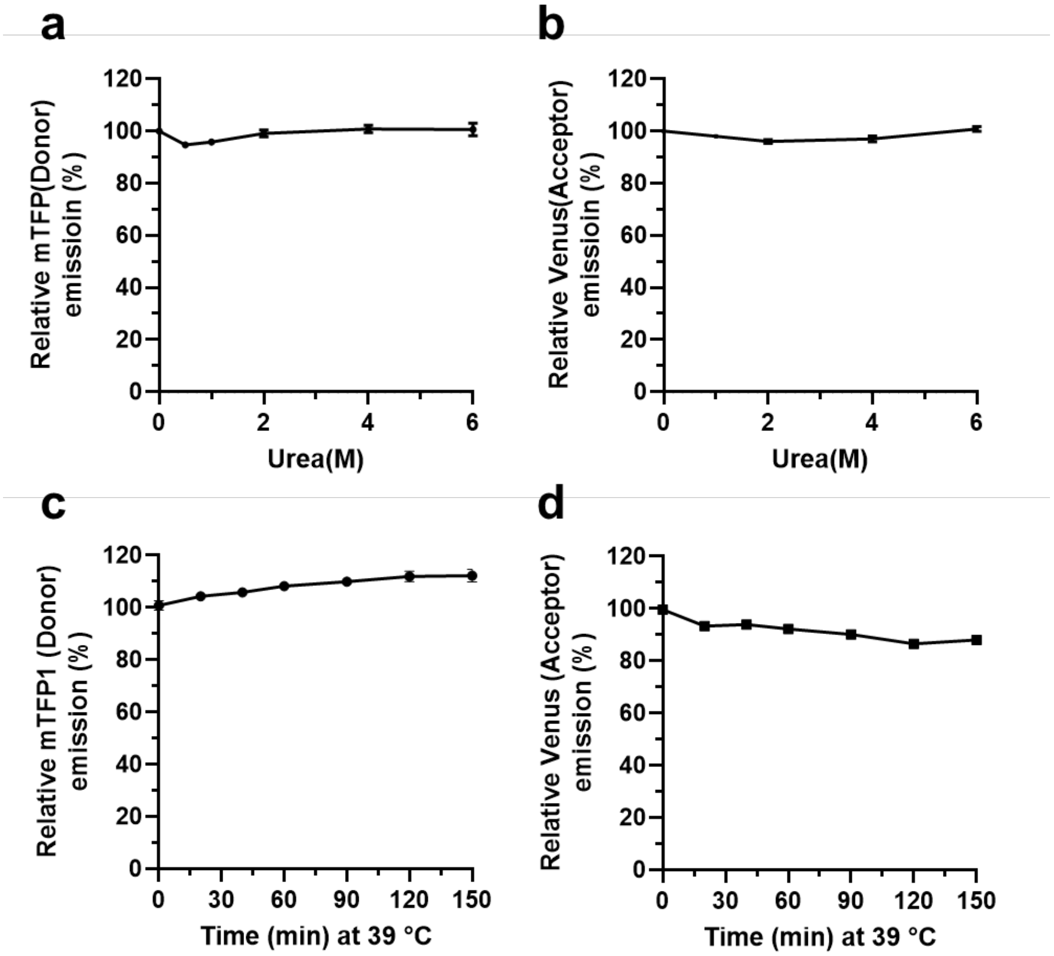


**Figure S2:** Urea and temperature stability of individual mTFP1 (left) and Venus (right). The intrinsic fluorescence of 0.4 μM mTFP1 (**a**) or Venus (**b**) was measured at 30°C for 15 min in the presence of up to 6 M Urea and normalized to the no urea control (100%). The intrinsic fluorescence of 0.4 μM mTFP1 (**c**) or Venus (**d**) at 39°C was measured during up to 150 minutes and normalized to the fluorescence at 30°C at T=0 .

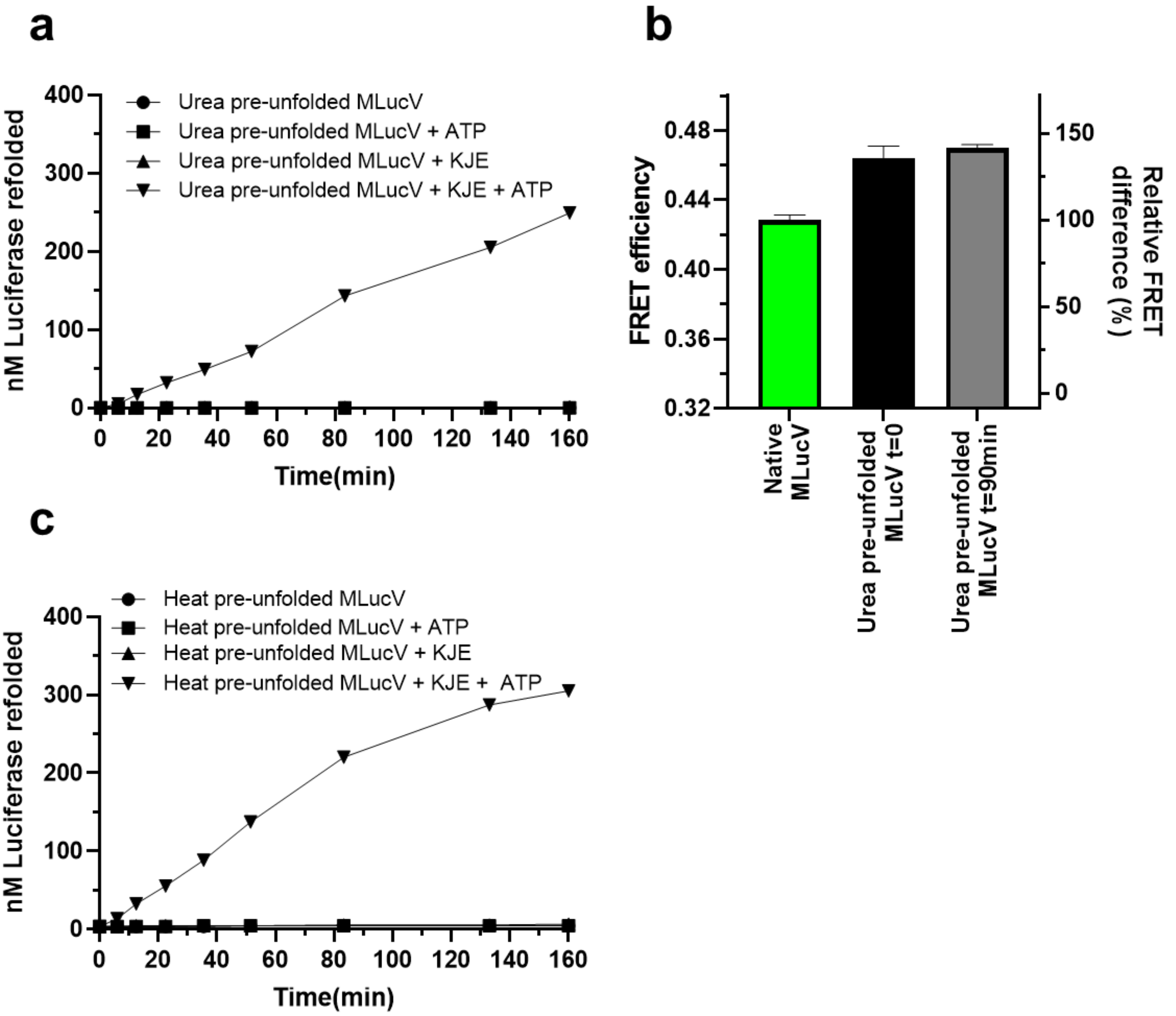


**Figure S3. a)** 30 μM MLucV was incubated 5 min in 4M urea and then diluted 75 folds into buffer containing 4 μM Bovine Serum Albumin (BSA) and either no ATP and no chaperones, or 4 mM ATP, of KJE (4:1:2 μM respectively). **b)** Calculation (performed as in Fig. 2b) of FRET efficiencies of urea pre-unfolded MLucV directly after dilution (t=0) and after 90min, compared to native MLucV. **c)** 400nm MLucV was incubated 32 min at 39.8°C in refolding buffer containing 4 *μ*M BSA and either no ATP and no chaperones, or 4 mM ATP, of KJE (4:1:2 *μ*M respectively); and then shifted to 25°C and incubated up to 160 min. Numbers indicate yield (in nM) of active Luciferase refolded.

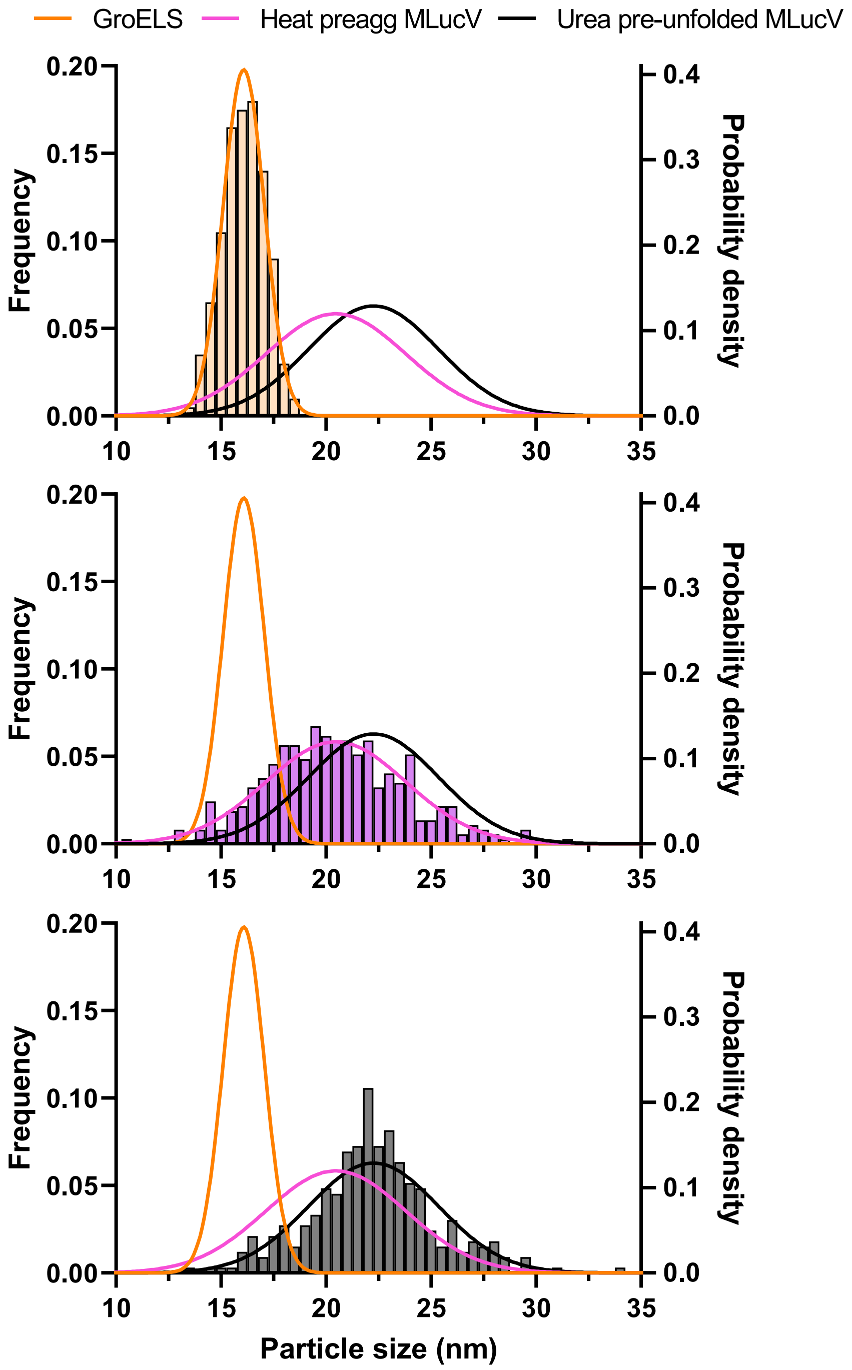


**Figure S4: Size distributions of particles observed by negative-staining TEM.** Particles sizes were manually measured using ImageJ. Histograms (left y-axes) of particle sizes are shown for GroEL rings (top, n=200), heat-preaggregated MLucV particles (middle, n=372), and aggregates from urea pre-unfolded MLucV (bottom, n=331). For comparisons, all normal distributions fitted to the histograms are shown in each panel (right y-axes).

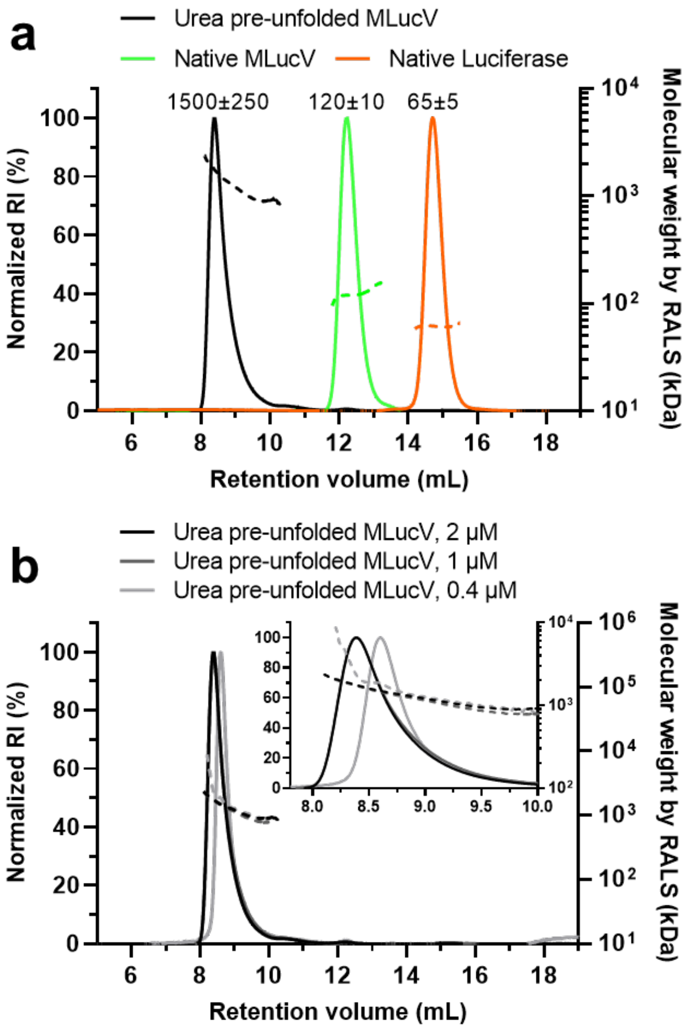


**Figure S5**. **Molecular weight calculations of native and aggregated MLucV by SEC-RALS**. In all panels, left y-axes show the relative abundance of eluted proteins, measured by differential refractive index (RI); and right y-axes show the calculated molecular weights (shown as dashed lines) of eluting species. **a)** SEC-RALS analysis of Native MLucV, native Luciferase monomers and urea pre-unfolded MLucV, showing elution of urea pre-unfolded MLucV as a ~12 mer (black, 1500 ± 250 kDa), native MLucV as a monomer (green, 120 ± 10 kDa), and native Luciferase as monomer (orange, 65 ± 5 kDa). **b)** SEC-RALS analysis of different concentrations of MLucV pre-unfolded in urea. The inset shows the narrow range of elution times and molecular weights.

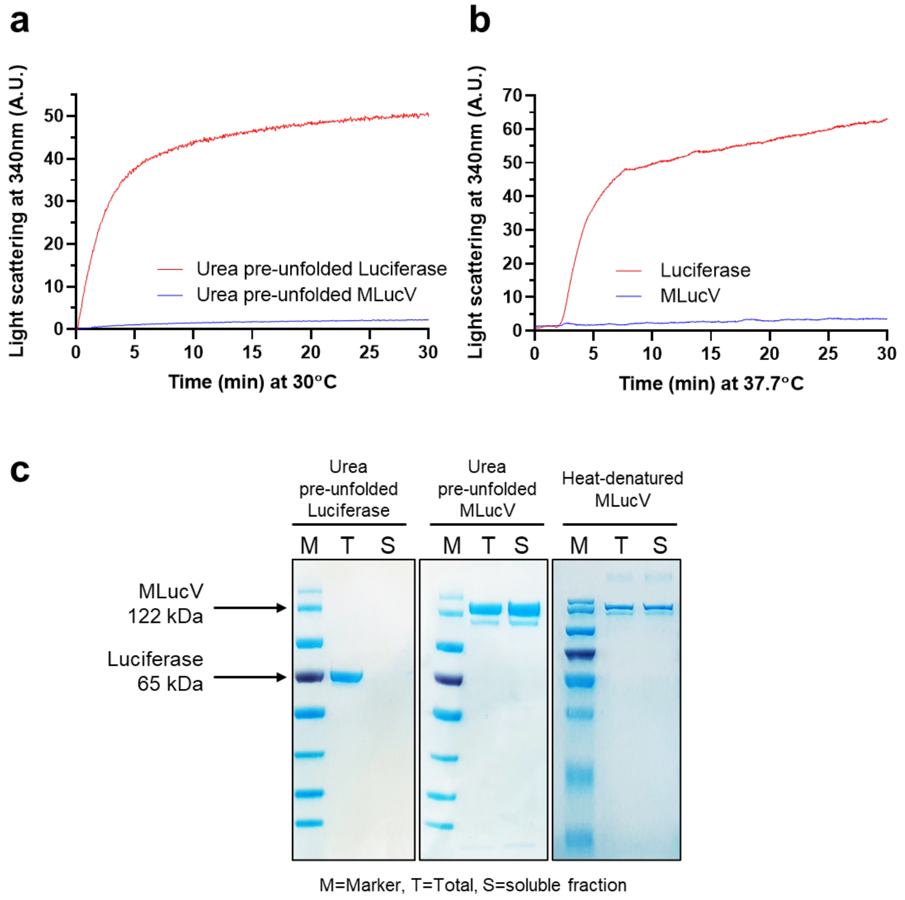


**Figure S6: Aggregation propensity of Urea pre-unfolded or heat-denatured Luciferase and MLucV. a**) 20 $\mu$M Luciferase or MLucV was preunfolded with 4 M urea at 25°C for 30 min, then diluted to a final concentration of 1 $\mu$M in buffer (50mM HEPES-KOH pH 7.5,150mM KCl, 10mM Mgcl2,2mM DTT), immediately aggregation was monitored by light scattering at 340 nm at 30°C for 30 min using Perkin Elmer Fluorescence Spectrophotometer. **b**) Aggregation propensity of heat denatured Luciferase and MLucV. 0.5 $\mu$M MLucV or Luciferase was incubated at 37.7℃ and aggregation was monitored for 30 minutes by light scattering at 340 nm. **c**) Solubility analysis of urea pre-unfolded and heat denatured - MLucV, compared to luciferase alone. ΔLuciferase or MLucV was denatured with 4M urea at 30℃ for 30 min, then diluted to a final concentration of 1 $\mu$M in buffer (50 mM HEPES-KOH, pH 7.5+150mM KCl+10mM Mgcl2+2mM DTT). Samples were incubated at 25°C for 30 min then the soluble fraction was separated with high-speed centrifugation (14000 rpm, 10 min). Equal volumes of Total and supernatant was loaded in 12% SDS-PAGE blue gel. For solubility after heat-denaturation, 0.5 $\mu$M MLucV was incubated at 37.7°C for 30 min then the soluble fraction was separated with high-speed centrifugation (14000 rpm, 10 min). Equal volumes of Total and supernatant was loaded in 12% SDS-PAGE gel stained with Coomassie.

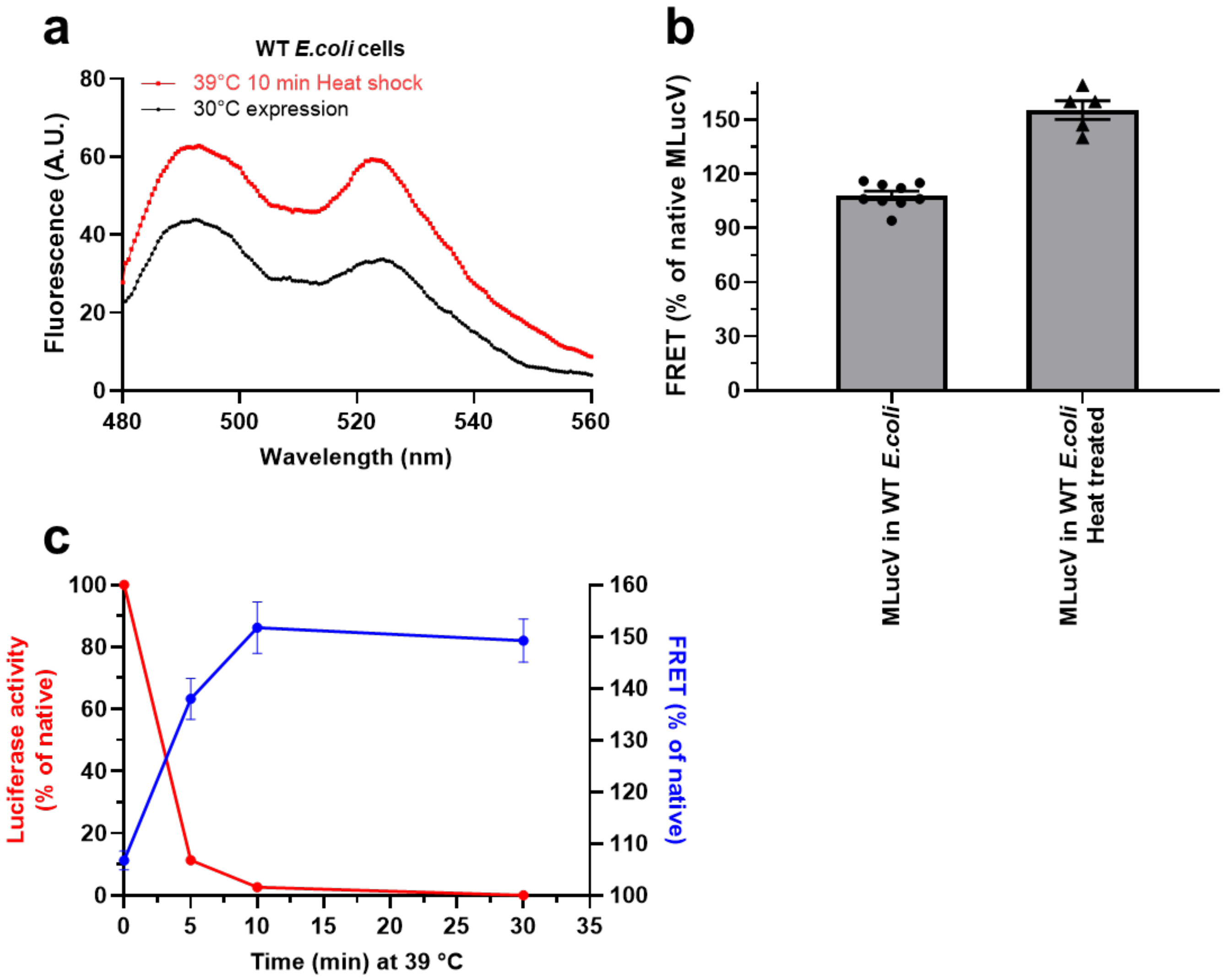


**Figure S7: Using MLucV as an *in-vivo* misfolding sensor**. **a)** FRET spectra of MLucV expressed under limited-growth conditions in WT W3110 *E. coli*; under native conditions at 30°C (black), or after a 10-min heat shock at 39°C (red). **b)** Calculation of FRET efficiencies from the spectra in a), normalized to that of native purified MLucV, with at least six biological repeats. **c)** Time course of MLucV misfolding in WT W3110 *E. coli* (see Methods) during heat shock at 39°C, measured by FRET (blue), and enzymatic activity (red).
